# A comparative analysis across species of maternal mRNA regulation in embryos using QUANTA

**DOI:** 10.1101/2024.06.27.600954

**Authors:** Mazal Tawil, Dina Alcalay, Pnina Greenberg, Shirel Har-Sheffer, Lior Fishman, Michal Rabani

## Abstract

The post-transcriptional regulation of mRNAs greatly impacts gene expression dynamics, but the underlying regulatory kinetics and sequence rules and how they change between organisms remain elusive. Thousands of pre-loaded maternal transcripts are post-transcriptionally regulated within metazoan embryos, making it an ideal system to investigate mRNA regulation. We present QUANTA, a computational strategy to distinguish transcriptionally silent genes and analyze their regulation. QUANTA uses kinetic models to compare total and polyA+ expression patterns, and dissect quantitative rates of mRNA polyadenylation and degradation. QUANTA analysis of maternally provided mRNAs in zebrafish, frog, mouse and human embryos shows that widespread polyadenylation precedes their degradation. Degradation rates are proportional to the developmental pace of organisms and diverge between orthologs. Rates also scale by adjusting developmental pace of zebrafish with external temperature. Finally, we implement a massively parallel reporter assay that is compatible with QUANTA analysis in zebrafish embryos, and analyze the effects of 3’UTR sequences on mRNA kinetics. We pinpoint potential regulatory signals in 3’UTRs of each organism. These reveal signals to accelerate maternal degradation in fast-developing organisms, while in slow-developing organisms signals enhance mRNA stability. Our work provides a general strategy to quantify post-transcriptional mRNA kinetics and investigate its sequence-based rules.

## Introduction

mRNA turnover is a critical determinant of gene expression levels within cells. Differences in turnover rates between mRNAs can span a few orders of magnitude (Fishman et al., 2024; Rabani et al., 2014) and significantly affect gene expression. For example, many housekeeping genes have low turnover, allowing cells to save energy and resources. On the other hand, high turnover is more expensive but allows for faster responses, thus characterizes stress-induced genes and cell identity factors (Li et al., 2023; Pérez-Ortín et al., 2007; Rabani et al., 2011; Yang et al., 2003). The stability of mRNAs also adapts in response to various stimuli, and facilitate rapid changes in gene expression (Choi et al., 2007; Rabani et al., 2014; Shalem et al., 2008; Viegas et al., 2022). For example, in mouse immune dendritic cells, degradation of key immune regulators increases during a pathogen response, allowing for their precisely timed expression (Rabani et al., 2014). During embryogenesis, maternally provided mRNAs are destabilized and cleared from embryos, enabling newly synthesized zygotic mRNAs to establish distinct developmental programs (Jukam et al., 2017). Similarly, during gastrulation, the destruction of Nodal agonist and antagonist mRNAs generates the precise expression levels needed for correct patterning (Choi et al., 2007).

The stability of mRNAs is regulated by interactions between cis-acting elements encoded within mRNA sequences, and trans-regulators that bind them. Regulators such as RNA binding proteins (RBPs) (Guo et al., 2017; Piqué et al., 2008; Ray et al., 2013; Siddall et al., 2006; Wharton and Struhl, 1991) and micro-RNAs (miRNAs) (Giraldez et al., 2006; Guo et al., 2010) selectively bind specific mRNAs and affect their fate. For example, the zebrafish miRNA miR- 430 (Giraldez et al., 2006) or the fly SMAUG RBP (Tadros et al., 2007) control the degradation of maternal mRNAs in embryos. Regulation is often also mediated by post-transcriptional modifications of mRNA molecules. In particular, the post-transcriptionally added poly(A) tail mediates changes to both mRNA stability and translation. Studies show that different mRNAs maintain different poly(A) tail lengths (Chang et al., 2018; Eichhorn et al., 2016; Lim et al., 2016; Subtelny et al., 2014). The binding of different regulators mediate the shortening or elongation of poly(A) tails (Passmore and Coller, 2022), thereby directing or protecting mRNAs from degradation. For example, tightly regulated changes of maternal poly(A) tails accompany oocyte maturation and early embryogenesis in many organisms (Chang et al., 2018; Eichhorn et al., 2016; Lim et al., 2016; Subtelny et al., 2014).

Despite their central role in regulating gene expression, technical and biological difficulties limit our ability to systematically measure mRNA stability and polyadenylation. Standard RNA-Seq approaches measure overall mRNA levels, but cannot distinguish the degradation of pre-existing mRNAs from the accumulation of new transcripts. To address this challenge, RNA-Seq is combined with strategies for transcriptional arrest (Lai et al., 2019; Viegas et al., 2022) or RNA metabolic labeling (Lugowski et al., 2018; Rabani et al., 2011; Russo et al., 2017; Tani and Akimitsu, 2012). Such approaches utilize manipulations of the biological system that might affect their accuracy. Alternatively, quantification of intron-containing pre-mRNAs (Furlan et al., 2020; Lee et al., 2013) and single nucleotide polymorphisms (SNPs) (Harvey et al., 2013) provide native information for such analyses. Nonetheless, interpretation of degradation rates from such signals is indirect, and could be affected by changes in regulation, such as splicing rates (Rabani et al., 2014). In addition, a main challenge in studying poly(A) tails is that they are difficult to accurately measure (Nicholson and Pasquinelli, 2019). Advanced analysis technologies of sequencing data has enabled a genome-wide look at poly(A) tails by RNA-Seq (Chang et al., 2018; Krause et al., 2019; Subtelny et al., 2014). However, their application remains limited by the accessibility of the needed sequencing adaptations. Alternatively, comparing gene expression levels between polyA+ and total-RNA fractions was also shown effective to indirectly estimate poly(A) tail lengths (Slobodin et al., 2020; Winata et al., 2018). Still, such approaches were not combined with a quantitative analysis. As a result, tools for an accurate quantification of mRNA regulation from standard RNA-Seq measurements are still needed.

It has also been difficult to determine the sequence-based rules of mRNA stability. Prediction of cis-regulatory elements from genome-wide measurements is limited by the ability to distinguish and quantify regulatory processes on a genome-wide scale (Alonso, 2012; Hennig and Sattler, 2015; Kim and Wysocka, 2023; Medina-Muñoz et al., 2021). In addition, prediction of binding sites commonly relies on a discrete set of positive and negative examples (Bailey, 2021; Bailey et al., 2015). Such tools are not well-suited to analyze complex outcomes such as mRNA stability, which span a wide range of activities across genes due to combinatorial effects of multiple regulators. To mitigate those complexities, large-scale reporter assays have been used to identify sequence elements that affect mRNA stability (Rabani et al., 2017; Vejnar et al., 2019) and polyadenylation (Xiang et al., 2024). But application of their results to native transcripts is limited by multiple competing effects and combinatorial interactions, which are not well captured by the reporter approach (Medina-Muñoz et al., 2021). Thus, our ability to predict genome-encoded cis-regulatory elements and associate them with quantitative mRNA regulation is still limited.

The massive degradation of maternal mRNAs is a key regulatory event in early metazoan embryos (Vastenhouw et al., 2019) and a powerful system to study mRNA degradation in the absence of de-novo transcription. At the onset of development, embryos are transcriptionally silent, and utilize maternally provided mRNAs and proteins for all functionality. As development progresses, thousands of maternally provided transcripts are rapidly degraded and replaced with newly synthesized zygotic mRNAs. The zebrafish maternal-to-zygotic transition is a well-studied example in vertebrates. Three hours post fertilization (hpf) zebrafish newly synthesized zygotic mRNAs start to gradually replace destabilized maternal transcripts and establish distinct zygotically encoded developmental programs (Jukam et al., 2017). Several pathways for zebrafish maternal mRNA degradation have been documented after genome activation, including miR-430 mediated regulation (Giraldez et al., 2006), inefficient codon compositions (Bazzini et al., 2016; Mishima and Tomari, 2016) and lower levels of m^6^A (Zhao et al., 2017) and m^5^C (Yang et al., 2019) modified bases. Control of maternal transcripts prior to genome activation tightly relies on changes in their poly(A) tail (Chang et al., 2018; Subtelny et al., 2014), with contradicting evidence that either implicated (Aanes et al., 2011; Voeltz and Steitz, 1998) or ruled out early mRNA decay (Vesterlund et al., 2011). Unlike fast-developing anamniotic embryos (e.g., fish and amphibians), amniotes embryos (e.g., mammalian) develop more slowly (Jukam et al., 2017). These two vertebrate developmental modes have both conserved principles and important differences in maternal mRNA regulation (Jukam et al., 2017; Shen-Orr et al., 2010). For example, RBP-mediated degradation (Ramos et al., 2004; Yu et al., 2016; Zheng et al., 2020) and lower levels of m^5^C modified bases (Liu et al., 2022) are shared in mammalian maternal mRNA degradation, but miRNA-mediated repression is not active (Suh et al., 2010). Thus, despite important advances in our understanding of the vertebrate maternal-to-zygotic transition, it is still unclear how maternal mRNA degradation patterns change between organisms, and if similar regulatory rules apply.

Here we develop QUANTA (QUantitative ANalysis of Total and A+ RNA), a kinetic modeling framework to analyze interconnected mRNA polyadenylation and degradation programs of transcriptionally silent genes within temporal RNA-Seq datasets. We build and validate QUANTA using the well-studied example of the zebrafish maternal-to-zygotic program. We combine QUANTA with a massively parallel reporter assay that measure both polyA+ and total-RNA in zebrafish embryos, and associate differences in regulation with putative cis-regulatory signals. Expanding our analysis to frog, mouse and human embryogenesis, we determine conserved and divergent patterns of maternal mRNA regulation and cis-elements between organisms, and demonstrate its broad utility. QUANTA is a general strategy to analyze mRNA degradation programs and investigate their sequence-based rules across species and biological systems.

## Results

### Dissecting the maternal and zygotic transcriptomes by pre-mRNA expression

To identify actively transcribed genes, QUANTA analyzes (see **Methods**) the presence of pre-mRNA precursors in total-RNA-Seq data from zebrafish embryos (**Fig. 1A**). For each gene, we define a precursor isoform (pre-mRNA) that spans the entire gene locus including intronic regions, and quantify pre-mRNA in parallel to other mRNA isoforms by calculating isoform-specific mRNA abundance (Li and Dewey, 2011). Actively transcribed genes, such as zygotically expressed embryonic transcripts (e.g., *fgfr4*, **Fig. 1B**), are co-transcriptionally processed and spliced and thus their introns are also detected after genome activation. On the other hand, maternally provided mRNAs (e.g., *lhx8a*, **Fig. 1B**) are pre-deposited into the egg and present in the embryo as fully spliced mature transcripts that do not contain introns. Some genes are both pre-deposited and expressed (e.g., *tubb4b*, **Fig. 1B**), thus introns are not detected initially, but start to appear after genome activation. Therefore, we identify new transcription by an increase in expression levels of either mRNA or pre-mRNA isoforms after genome activation in comparison to its earlier levels (**Methods**). Only mRNAs which do not exhibit new transcription are classified as maternal. Zygotic genes, on the other hand, lack expression prior to genome activation.

**Figure 1.**
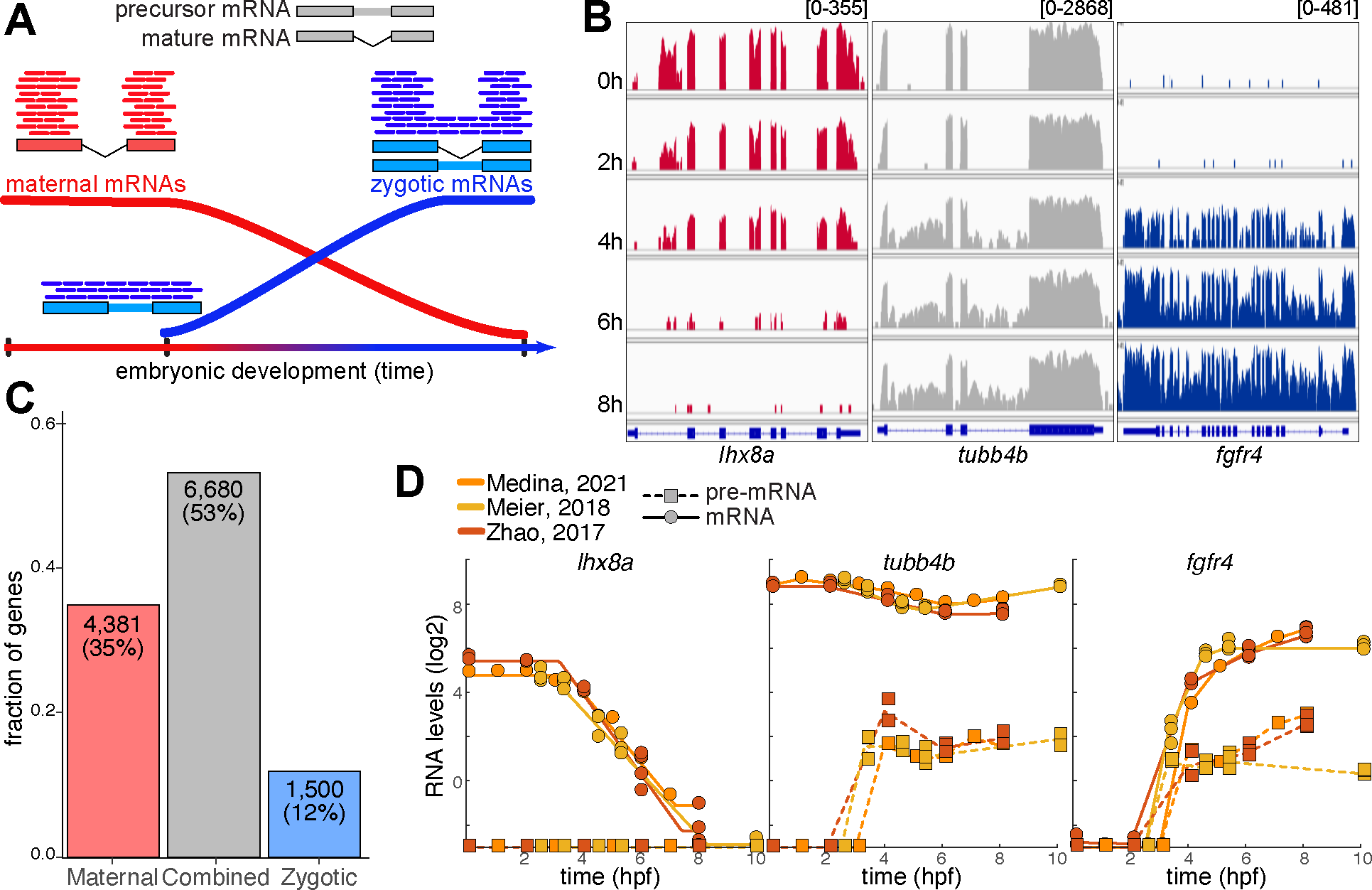
Dissecting the maternal and zygotic transcriptomes by unspliced mRNA expression **(A)** Early animal development is characterized by transcriptionally silent embryos that utilize maternally provided mRNAs (red). These are present in the embryo as fully spliced and processed transcripts, with total-RNA-Seq reads (lines) that match only exon sequences (boxes). As development proceeds (x-axis, time), maternal transcripts are degraded and replaced with newly synthesized zygotic mRNAs (blue). Zygotic mRNAs are actively transcribed and therefore also processed and spliced by the embryo. As a result, their intron sequences (lines connecting boxes) are also detected in total-RNA-Seq. **(B)** Top: total-RNA-Seq coverage (y-axis, scale of coverage is noted on top) in temporal samples (hours, top to bottom) of zebrafish development. Bottom: gene name and structure (exons: box, introns: line). Red (left): a maternal gene (*lhx8a*). Coverage is only on exons. Blue (right): a zygotic gene (*fgfr4*). Expression only after zebrafish genome activation (∼3.5 hpf), and coverage spans both exons and introns. Gray (middle): combined expression (*tubb4b*). Before genome activation, coverage is only of exons before genome activations, and introns are covered after genome activation. **(C)** Classification (x-axis: class) of zebrafish embryonic genes (y-axis, fraction of genes). Number of genes and their factions are indicated on bars. **(D)** Temporal (x-axis, hpf) total-RNA-Seq expression levels (y-axis, normalized FPKM, log2) in three analyzed datasets of unspliced pre-mRNA (dashed/square) and fully spliced mRNA (solid/circle) for the three example genes shown in (B).

This approach successfully distinguishes maternally provided from zygotically expressed or combined (maternal and zygotic) embryonic transcripts. We validate our approach by applying it to three published total-RNA-Seq datasets of zebrafish embryos during the first 10 hours of development (Medina-Muñoz et al., 2021; Meier et al., 2018; Zhao et al., 2017). First, we quantify minimal pre-mRNA levels prior to zygotic genome activation, and a substantial increase in samples that were collected after activation (**Fig. S1**). Second, looking at canonical maternal or zygotic genes, we find that only maternal mRNAs are detected at early times (**Fig. S2A**). On the other hand, an increase in both mRNA and pre-mRNA levels is characteristic of zygotic genes (**Fig. S2B**). Third, comparing our classification between datasets, we consistently classify 70% of genes (8,803/12,561) in all 3 datasets, and additional 29% (3,673/12,561) in 2 out of 3 datasets (**Fig. S3A**). Our classification is also consistent with classification by two metabolic labeling datasets (Bhat et al., 2023; Fishman et al., 2024), for 56% of genes in all 3 datasets, and 36% of genes with at least one of the two metabolic labeling datasets (**Fig. S3B**).

Our final classification (**Table S1**) is based on the majority classification between the 3 total-RNA-seq datasets above for each gene (**Fig. 1C**). In line with previous reports (Aanes et al., 2011; Holler et al., 2021; Rabani et al., 2014; White et al., 2017), we find that most embryonic genes are both maternally provided and zygotically expressed. We classify 35% (4,381/12,561, e.g., *lhx8a*, **Fig. 1D**) genes as maternal only, another 12% (1,500/12,561, e.g., *fgfr4*, **Fig. 1D**) genes as zygotic only and 53% (6,680/12,543, e.g., *tubb4b*, **Fig. 1D**) genes as combined (maternal and zygotic) expressed. Functional annotations within each group match expected biological processes, and further validate our classification (**Table S2**). Maternal genes are enriched for reproductive process (5% FDR hypergeometric p<3*10^-6^), sperm-egg recognition (p<4*10^-5^) and oogenesis (p<5*10^-4^). On the other hand, zygotic genes are enriched for developmental process (p<2*10^-23^), tissue development (p<2*10^-17^) and regionalization (p<5*10^-17^). Finally, combined maternal and zygotic genes are enriched for many housekeeping processes such as mRNA splicing (p<3*10^-46^), translation (p<7*10^-49^) and chromatin organization (p<2*10^-52^).

Taken together, these results validate our approach for identifying actively transcribed genes by pre-mRNA expression. Its application classifies embryonic genes as maternal or zygotic. This approach can be easily applied to any total-RNA-Seq time course dataset, and does not require any manipulations that might interfere with normal development and growth.

### Early changes in polyA+ maternal RNAs are not paralleled in total RNA

Expression patterns of maternally deposited genes that are not transcribed by the embryo reflect regulation by mRNA degradation, and allow a direct quantitative view on the kinetics of mRNA decay. We analyzed the temporal expression patterns of 4,381 zebrafish maternal genes in the 3 total-RNA-Seq datasets we used for classification, and in additional 4 published polyA+ RNA-Seq datasets of zebrafish embryos (Harvey et al., 2013; Medina-Muñoz et al., 2021; Pauli et al., 2012; Yang et al., 2019). These additional datasets measure only the polyA+ mRNA fraction within samples, by applying poly(A) selection during library preparation.

Our analysis reveals marked differences in expression patterns between polyA+ and total-RNA populations. For many genes, the changes in expression observed in polyA+ datasets are not paralleled in total-RNA datasets (**Fig. 2A**). In particular, while total-RNA levels of most genes remain constant prior to zygotic genome activation (before 3 hpf), polyA+ levels of genes changes over time in corresponding measurement. For example, polyA+ levels of genes such as *mcm3l* and *buc* (**Fig. 2B**) increase before 3 hpf, and those of genes such as *wee2* and *dazl* (**Fig. 2B**) decrease before 3 hpf. However, at the same time, their total-RNA levels remain unchanged. Only few genes (e.g., *gdf3*, **Fig. 2B**) behave similarly in both populations. Globally, initial expression level of maternal genes (at 0-1 hpf) are on average 2.5-fold lower in polyA+ compared to total-RNA (**Fig. S3C**), and 48% of polyA+ RNA levels increase over time (>1.2- fold increase relative to the initial measurements taken at 0-1 hpf, **Fig. S3D**). On the other hand, 51% of total-RNA levels do not change over time (<1.2-fold decrease **Fig. S3D**). A notable fraction of measurements decreases over time in both populations (34% of polyA+ and 41% of total-RNA >1.2-fold decrease, **Fig. S3D**), as expected.

**Figure 2.**
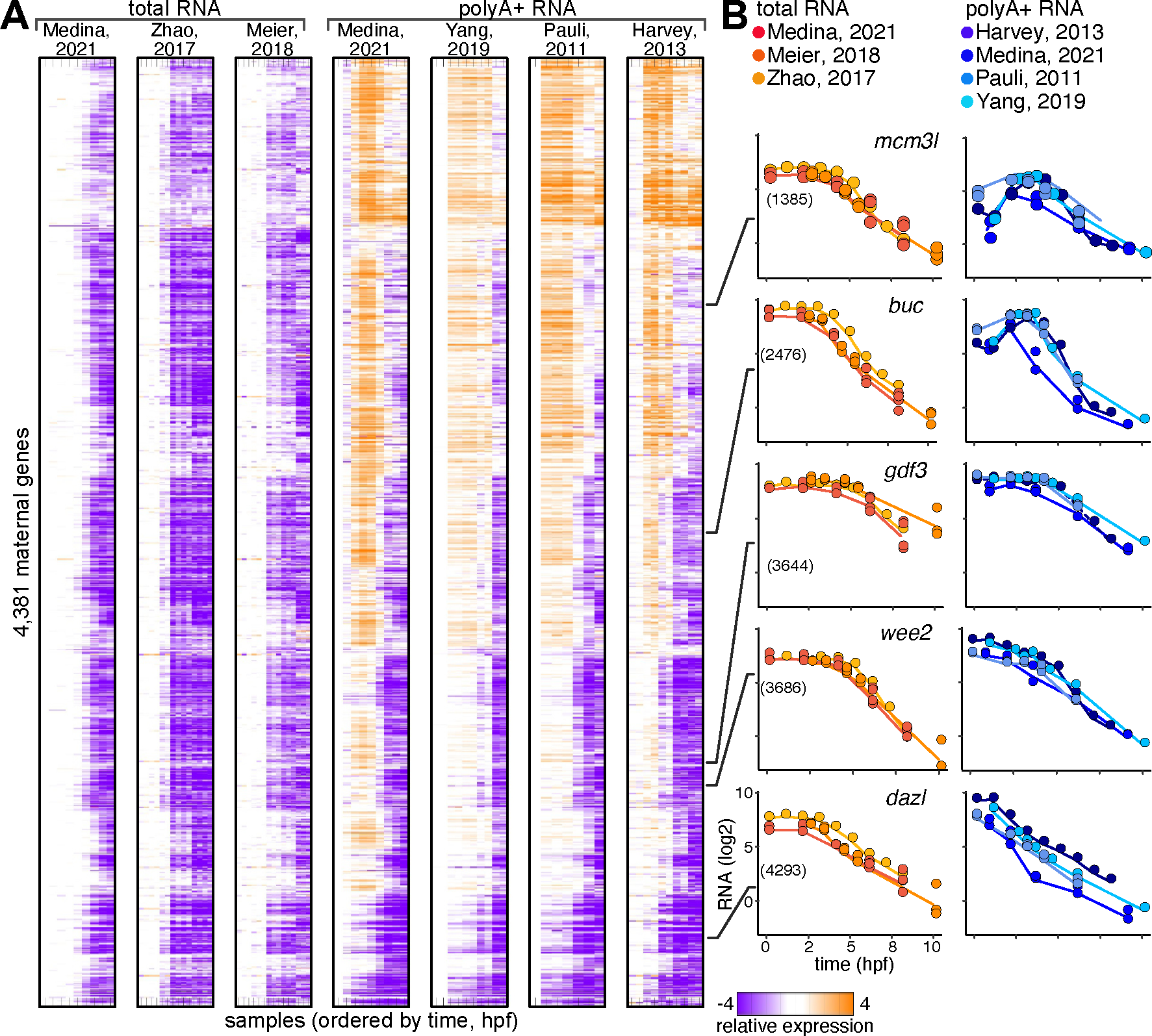
The temporal dynamics of zebrafish maternal mRNAs **(A)** Temporal (columns) mRNA expression patterns relative to initial measurement (orange: increase; purple: decrease; white: unchanged) of 4,381 zebrafish maternal genes (rows) in 3 total-RNA-Seq datasets (left), and 4 polyA+ RNA-Seq datasets (right) of early zebrafish embryogenesis. **(B)** Examples of temporal (x-axis, hpf) expression levels (y-axis, normalized FPKM, log2) measured in 3 total-RNA-Seq datasets (left, orange) and 4 polyA+ RNA-Seq datasets (right, blue) for specific maternal genes (names are indicated on graphs).

These observations identify changes in polyA+ expression of maternal genes that are not paralleled in total-RNA expression. As there is no transcription of new mRNAs prior to genome activation, we suggest that these differences reflect changes in the average poly(A) tail length of mRNAs, which influence their recovery in the polyA+ fraction (Park et al., 2016; Winata et al., 2018). A longer average poly(A) tail length may allow better recovery.

### Kinetic models of total and polyA+ maternal RNA expression capture in-vivo dynamics

To investigate this hypothesis, we implemented in QUANTA a kinetic modeling scheme that uses two alternative models for the dynamics of maternal mRNAs degradation (**Fig. 3A**). First, a simpler ‘degradation model’ assumes that levels of maternal mRNA are determined by an exponential decay of pre-existing copies. We assume that gene-specific degradation rates are constant over time, but incorporate a gene-specific onset-time and offset-time for degradation. An alternative ‘polyA+ model’, uses an additional rate constant to model early changes in expression levels until degradation onset. Early rate constant can be either positive to represent a decrease in mRNA levels (as in the “degradation model”), or negative and represent an increase in mRNA levels.

**Figure 3.**
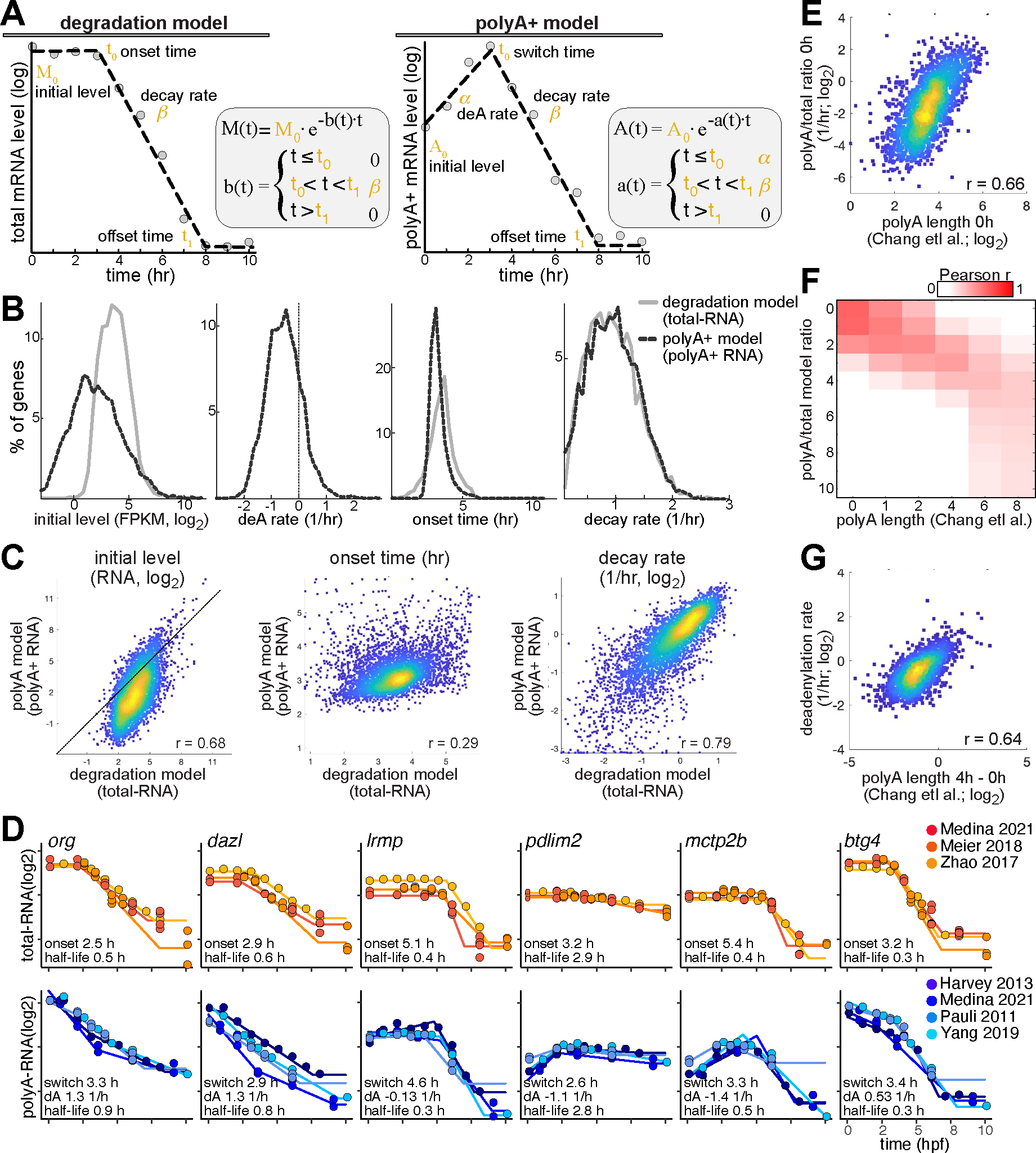
Kinetic models of total and polyA+ maternal gene expression elicit their regulatory kinetics **(A)** Two alternative models for the dynamics of maternal mRNA degradation. Both models model temporal changes in maternal RNA levels by exponential changes of pre-existing copies, but use different rate functions. Positive rates are associated with an exponential decrease in mRNA levels, while negative rates are associated with an exponential increase. Left: a “degradation model” for analyzing maternal total-RNA levels (M0, initial level). A gene-specific rate function is parametrized by a constant decay rate (β, 1/hr), an onset-time (t0) and an offset-time (t1). Right: a “polyA+ model” models for analyzing maternal polyA+ RNA levels (A0, initial level). A gene-specific rate function is parametrized by two constant rates: an early deadenylation rate (α, 1/hr) and a late decay rate (β, 1/hr) with a switch-time (t0) and an offset-time (t1). **(B)** Distribution (y-axis, % of maternal genes) of per-gene average of parameter values (x-axis) estimated by models in embryonic RNA-Seq datasets (gray: degradation model, black: polyA+ model). Left to right: initial RNA level (normalized FPKM, log2), deadnylation rate (1/hr), onset/switch time (hr) and decay rate (1/hr). **(C)** Correlation between average parameter values predicted by the “degradation model” on total-RNA-Seq datasets (x-axis), and “polyA+ model” on polyA+RNA-Seq datasets (y-axis). Left to right: initial RNA level (normalized FPKM, log2), onset/switch time (hr) and decay rate (1/hr, log2). Color represents density (blue = low density; yellow = high density). **(D)** Fitting of models to temporal (x-axis, hpf) total (top, orange) and polyA+ (bottom, blue) expression levels (y-axis, normalized FPKM, log2) of specific maternal genes in all analyzed datasets. Gene names and average parameters of the fitted models are indicated. **(E)** Correlation between polyA tail length measured at 0 hpf by Chang et. al. (x-axis, log2) and the ratio of estimated polyA+ and total RNA levels at 0 hpf by our models (y-axis). Pearson r value is indicated. Color represents density (blue = low density; yellow = high density). **(F)** Heatmap representing Pearson r values for comparison between polyA tail lengths measured by Chang et. al. (x-axis, log2) at different times and the ratio of estimated polyA+ and total RNA levels by our models (y-axis) at different times. **(G)** Correlation between difference in polyA tail lengths measured at 4hpf and 0hpf as by Chang et. al. (x-axis, log2) and average deadenylation rates predicted by “polyA+ model” on polyA+-RNA-Seq datasets (y-axis, 1/hr, log2). Pearson r value is indicated. Color represents density (blue = low density; yellow = high density).

To validate our modeling approach, we determine the ability of the two models to capture temporal expression changes in maternal mRNAs. We apply the models to quantify the kinetics of 4,381 zebrafish maternal genes in each of 3 total-RNA-Seq and 4 polyA+ RNA-Seq datasets, after validating noise limits of each dataset by simulation studies (**Fig. S4A**). For each gene, we find the parameters that best fits the temporal measurements (**Methods**) by each of the two models. Both models capture a large fraction of the variability in the expression data. In 3 total-RNA-Seq datasets, either the ‘degradation’ or ‘polyA+’ models capture 90-98% (r-squared) of expression variability across maternal genes (**Fig. S4B**). In 4 polyA+ RNA-Seq datasets, the simpler ‘degradation model’ captures 87-93% of expression variability, and the ‘polyA+ model’ improves this by 5-9% (**Fig. S4B**).

To further test the suitability of the two models to different datasets, we also compare them via a likelihood ratio test. In total-RNA-Seq datasets, less than 17% of genes reject the ‘degradation model’ in favor of the ‘polyA+ model’ alternative (**Fig. S4C**), compared to 75-90% of genes in polyA+-RNA-Seq datasets (**Fig. S4D**). Indeed, the simpler ‘degradation model’ fits 92% of genes with an r^2^ > 0.7 in 3 total-RNA-Seq datasets (**Fig. S4E**). Application of the ‘polyA+ model’ to these datasets does not provide any significant improvement. However, in 4 polyA+ RNA-Seq datasets, 93% of genes has an r^2^ > 0.7 with the ‘polyA+ model’, compared to only 71% by the ‘degradation model’ (**Fig. S4E**). Finally, we also validate that the kinetic parameters predicted by the models are consistent between datasets (**Fig. S5A-B**), demonstrating the reproducibility of our predictions.

Taken together, these results show that the QUANTA kinetic models successfully capture and quantify the regulatory kinetics of maternal transcripts from temporal RNA-Seq measurements. The simpler ‘degradation model’ accurately captures total-RNA-Seq data, while the ‘polyA+ model’ is better suited to model polyA+ RNA-Seq datasets. Analysis of total- and polyA+ RNA-Seq by QUANTA reliably and reproducibly quantify maternal mRNA kinetic parameters in-vivo.

### A quantitative analysis distinguishes two phases of cytoplasmic mRNA metabolism in embryos

Next, we utilized QUANTA’s predictions to investigate maternal mRNA metabolism in embryos. We analyze the kinetic parameters predicted by the ‘degradation’ and ‘polyA+’ models on total or polyA+ RNA-Seq datasets respectively, to establish differences and similarities in their dynamics. For this analysis, we use per-gene averaged kinetic parameters across polyA+ or total-RNA-Seq datasets (**Methods**), relying on demonstrated reproducibility.

The most notable difference between polyA+ and total-RNA dynamics are early changes in polyA+ RNA levels that are not paralleled in total-RNA. As expected, initial polyA+ RNA levels estimated by the models are 3.2-fold lower on average than total RNA levels (**Fig. 3B- C**). Subsequently, polyA+ RNA increases after fertilization for >72% of maternal transcripts (negative rates, e.g., *mctp2b* and *pdlim2*, **Fig. 3D**) and decreases in another 28% of genes (positive rates, e.g., *org* and *dazl*, **Fig. 3D**).

Since these changes precede genome activation, we hypothesize they reflect changes in the average poly(A) tail length of maternal transcripts, which are mediated by cytoplasmic polyadenylation and deadenylation of embryonic genes (Charlesworth et al., 2013; Winata et al., 2018). Indeed, the ratio of polyA+ to total RNA levels predicted by QUANTA (which we term “AT-ratio”) correlates to poly(A) tail lengths (Chang et al., 2018) measured in zebrafish embryos (**Fig. 3E-F**). Moreover, the early rate parameter predicted by the ‘polyA+ model’ correlates to differences in measured poly(A) tail length between 1 and 4 hpf (**Fig. 3G**). Therefore, we refer to this rate as “deadenylation rate”, which is estimated per gene. Thus, although prior to zygotic genome activation total-RNA levels of maternal transcripts remain constant, their average poly(A) tail length is extended and shortened.

Despite differences in the early phase of the response, both models predict comparable degradation rates after genome activation (**Fig. 3B-C**). This suggests that the massive maternal mRNA degradation after genome activation has similar rate in both polyA+ and total-RNA fractions. Overall, maternal half-lives range from less than 30 minutes to 4 hours and more (median of 45 min), similar to previous estimates in-vivo (Rabani et al., 2017, 2014). However, there are noticeable differences between maternal genes in their degradation rates (e.g., *btg4* 0.4 hr unstable, *pdlim2* 4.3 hr stable; **Fig. 3D**). Degradation of maternal transcripts onsets during a narrow 3-5 hpf time window, which overlaps zygotic genome activation (>90% of maternal genes; **Fig. 3C**). Thus, differences in onset time between genes are relatively small (e.g., *org*, *dazl* early onset < 3.5 hpf, *mctp2b, lrmp* late onset > 4.5 hpf; **Fig. 3D**). In mRNAs undergoing an initial increase in polyA+ RNA (poly(A) tail extension), degradation onsets earlier in polyA+ than in total-RNA (mean 3.1 vs. 3.5 hr, respectively). On the other hand, mRNAs undergoing an initial decrease in polyA+ RNA (poly(A) tail shortening), degradation onsets later in polyA+ than total-RNA (mean 3.5 vs. 3.1 hr, respectively). These differences suggest that deadenylation precedes degradation.

Changes in poly(A) tail length have been associated with mRNA degradation. Therefore, we also compare between degradation rates and AT-ratios or deadenylation rates predicted by QUANTA (**Fig. S5C**). Degradation rates are not correlated to deadenylation rates and AT-ratio before genome activation. However, degradation rates are correlated to AT-ratio at 6 hpf (Pearson r=0.35) and to the change in AT-ratio between 2.3 to 5.2 hpf (r=0.38). These suggest that changes in poly(A) tail length at the time window of degradation onset (between 2 to 5 hpf) are associated with how fast mRNAs are degraded.

Taken together, these observations distinguish two phases of cytoplasmic mRNA metabolism in embryos. Prior to genome activation most changes involve the poly(A) tail length, and are not correlated to degradation. Initially, most maternal transcripts have short poly(A) tails (Chang et al., 2018), which reduced their recovery in polyA+ fractions. These are extended or shortened while total mRNA levels remain constant. After genome activation, a massive mRNA degradation similarly affects both polyA+ and total-RNA.

### A massively parallel reporter assay recapitulates maternal mRNA regulatory kinetics in-vivo

To independently test these genomic observations, we implemented a massively parallel reporter assay (MPRA) approach that is compatible with QUANTA. In UTR-Seq, our MPRA approach (Rabani et al., 2017), we inject a large set of mRNA reporters with a constant backbone and different 3’UTR sequences into 1-cell stage embryos, and follow their mRNA levels over time. 3’UTR sequences of the reporters are all 110nt long and match annotated genomic 3’UTRs. In this new implementation, we use targeted RNA-Seq to follow the reporters both in total-RNA and in isolated polyA+ fraction (**Methods**, **Fig. 4A**). We assay two types of reporters, that were synthesized either with an initial 40nt poly(A) tail (A+), or without an initial tail (A-). Our sequencing depth allowed a reliable analysis of 2,186 different reporters.

**Figure 4.**
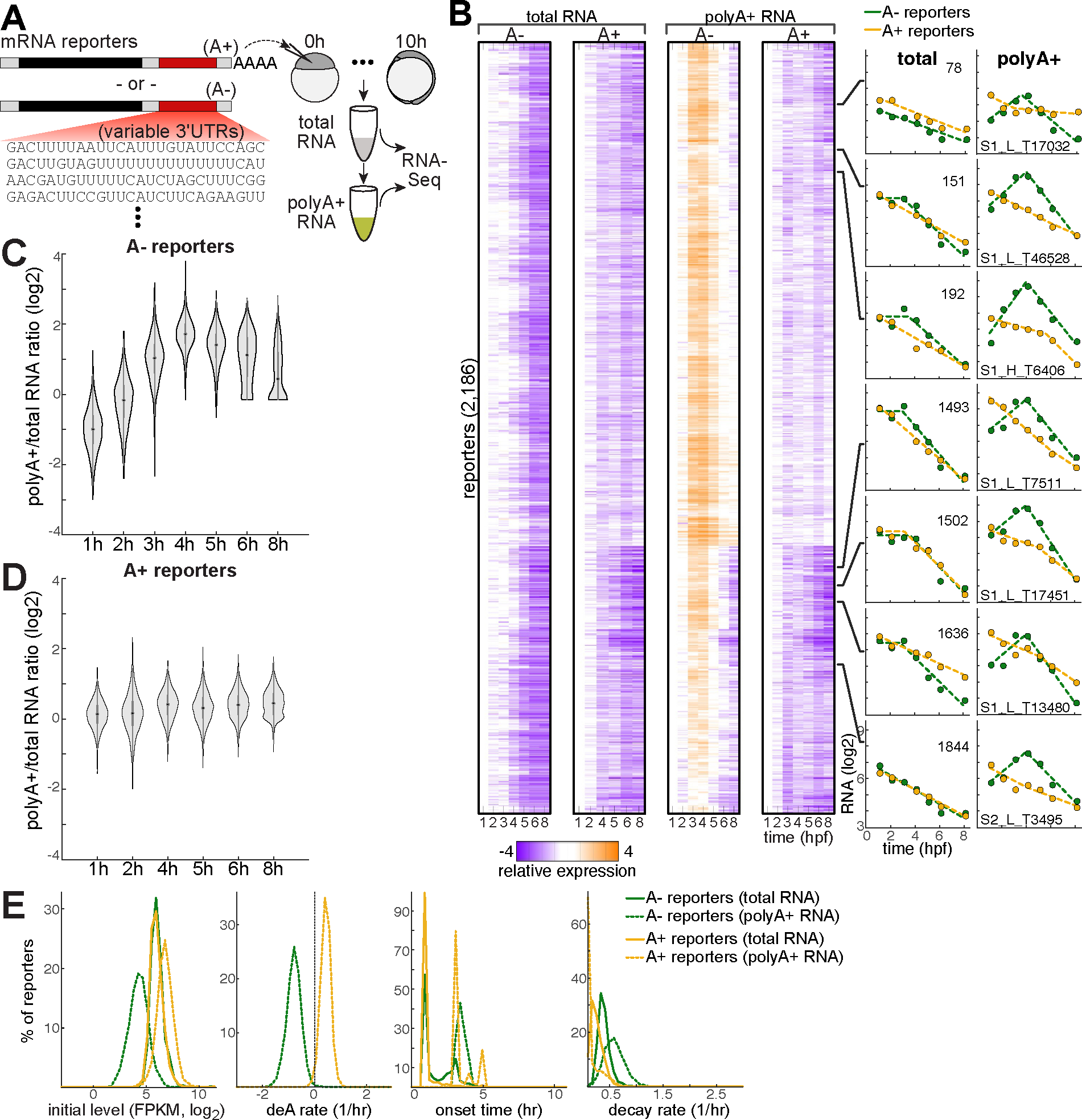
A massively parallel reporter assay recapitulates maternal mRNA regulatory kinetics in-vivo **(A)** A large set of mRNA reporters with a constant backbone and different 3’UTR sequences and polyA tail lengths were injected into 1-cell stage embryos and followed over time. Reporters were sequenced from temporal total- and polyA+ selected RNA samples. **(B)** Left: temporal (columns) mRNA expression patterns relative to initial measurement (orange: increase; purple: decrease; white: unchanged) of 2,186 mRNA reporters (rows) that were injected into zebrafish embryos at the 1-cell stage. Reporters were synthesized either with an initial polyA tail of 40nt (A+), or without an initial tail (A-) and captured in total- (left) or polyA+ (right) RNA fractions. Right: examples of temporal (x-axis, hpf) expression levels (y- axis, normalized FPKM, log2) of individual reporters (Left: total-RNA, right: polyA RNA; yellow: A+ reporters, green: A- reporters). **(C-D)** Distribution of measured polyA+ to total RNA level ratio (y-axis, log2) of reporters in temporal samples (x-axis). The central dot is median; gray box bounds are 25th and 75th percentiles, upper and lower limits of whiskers are 1.5x interquartile ranges. Values outside of the upper and lower limits are defined as outliers. **(C)** A- reporters. **(D)** A+ reporters. **(E)** Distribution (y-axis, % of reporters) of parameter values (x-axis) estimated by models in UTR-Seq datasets (yellow: A+ reporters, green: A- reporters; solid: degradation model, dashed: polyA+ model). Left to right: initial RNA level (normalized FPKM, log2), deadnylation rate (1/hr), onset/switch time (hr) and decay rate (1/hr).

Expression patterns of reporters recapitulate those of native maternal transcripts (**Fig. 4B**, **Fig. S6A**). First, initial (1 hpf) levels of A- reporters are on average 2-fold lower in polyA+ fraction compared to total-RNA, while levels of A+ reporters are similar (**Fig. S6B**), demonstrating dependence on poly(A) tail length in recovery. Second, total-RNA levels of reporters decrease over time, and most notably after genome activation. However, polyA+ RNA levels of A- reporters initially increase (compared to 1 hpf), and decrease after genome activation. Recapitulating the initial increase in polyA+ RNA levels with reporters rules out changes in expression as its source, since levels of these external constructs are set at injection and cannot increase further. Comparison between the two fractions further shows that polyA+ RNA levels of A- reporters increase relative to their total-RNA levels over time (**Fig. 4C**) but remain similar in both fractions for A+ reporters (**Fig. 4D**). These results therefore suggest that A- reporters become more abundant in polyA+ fraction due to extension of their poly(A) tails after their injection, while A+ reporters remain similarly present in both fractions.

QUANTA predictions reveal a repertoire of reporters’ regulatory kinetics, with similarities and distinctions from native maternal genes. Modeling analysis shows that for A- reporters, similar to native maternal genes, the ‘polyA+ model’ provides an improved fit to polyA+ RNA, but not to total-RNA (**Fig. S6C-E**). Interestingly, for A+ reporters, the ‘polyA+ model’ provides an improved fit to both polyA+ and total-RNA. These suggest that the kinetics of A+ reporters is different than native maternal genes, which generally have much shorter poly(A) tails at fertilization (Chang et al., 2018). These differences are also evident by several of the models’ parameters. First, predicted initial polyA+ RNA levels are lower than total-RNA levels for A- reporters as seen for most native maternal transcripts, but are higher for A+ reporters (**Fig. S6F**). Initial total-RNA levels are comparable between A- and A+ reporters (**Fig. 4E**). Second, similar to native maternal mRNAs, deadenylation rates of A- reporters are negative, representing a general increase in their polyA+ fraction due to an increase in their average poly(A) tail length (**Fig. 4E**). On the other hand, deadenylation rates of A+ reporters are positive, representing an early decrease in their polyA+ fraction. In this case, total-RNA levels also decrease in parallel, and thus this effect likely combines mRNA degradation with poly(A) tail shortening. These are distinguished by AT-ratios, which show different patterns between reporters (**Fig. 4D**). Third, unlike native maternal mRNAs, reporters’ degradation onset early in total-RNA, suggesting they are degraded also before genome activation (**Fig. 4E**). Only a small fraction of reporters (13% of A- reporters, 2% of A+ reporters) is stable until genome activation. Finally, degradation rates of reporters by polyA+ and total-RNA are correlated (**Fig. S6G**). However, A+ reporters are more stable than A- reporters (mean half-life by total-RNA 113 vs. 79 min., respectively), as previously shown (Rabani et al., 2017). Degradation of reporters is also slower on average than native maternal genes (mean half-life by total-RNA 45 min.).

Overall, these results validate our genomic observations on a set of synthetic mRNA reporters in-vivo. It recapitulates within a reporter system the early increase in polyA+ fraction, which is not paralleled by changes in total-RNA levels, and demonstrates that differences in poly(A) tail length affect recovery in polyA+ fraction. Subsequent decrease in maternal mRNA levels due to degradation occurs in both fractions at a similar rate. Results also highlight specific regulatory similarities and differences between injected reporters and native maternal transcripts, and suggest that A- reporters more closely recapitulate native maternal regulation.

### Decoding 3’UTR elements of the early and late phases of maternal mRNA regulation

Signals within 3’UTRs have been shown to regulate both polyadenylation and degradation of mRNA transcripts. Such signals are frequently interconnected and can affect both processes via multiple combinatorial regulators (Alonso, 2012). To investigate these signals, we analyzed 3’UTR sequences of maternal mRNAs with respect to the kinetic parameters quantified by our models. Using our previous approach (Fishman et al., 2024; Viegas et al., 2023), we systematically tested all 4-7 nucleotide long sequences (k-mers) within annotated 3’UTRs for their association with differences in parameters estimated by our kinetic models (**Methods**). We restricted our analysis both by significance (Kolmogorov-Smirnov FDR <1%) and effect size (standardized mean-difference).

These results identify both known and putative regulatory elements within 3’UTRs of maternal mRNAs (**Fig. 5A**, **Table S3**). Analysis of degradation rates confirms the destabilizing effect of the miR-430 seed sequence (GCACUU, p<5e-12), and AU-rich signals (ARE, e.g., AUAUUUA, p<2e-3). Destabilizing signals are also linked to lower AT-ratio after genome activation (GCACUU, p<1e-17 at 5hpf, AUAUUUA, p<8e-8 at 5hpf), suggesting shorter poly(A) tails, as expected. Nuclear polyadenylation elements (NPEs, e.g., AAUAAA) are associated with higher AT-ratios both before and after genome activation (p<8e-15 at 1hpf, p<1e-25 at 9hpf), and slower polyA+ degradation (p<5e-5), suggesting longer poly(A) tails. Several cytoplasmic polyadenylation elements (CPEs) are also identified (Charlesworth et al., 2013). Initially, CPEs are associated with higher AT-ratios and slower poly(A) tail extension rates, suggesting longer initial poly(A) tails. These include canonical CPEs (e.g., UUUUUA, p<1e-19 at 0hpf), eCPEs (e.g., UUUUUU, p<4e-10 at 1hpf) and Musashi binding elements (MBEs, e.g., UUUUAGU, p<3e-5 at 0hpf). After genome activation, CPEs are linked to delayed onset of total-RNA degradation and faster degradation rates, and switched their association to lower AT-ratios. Thus, CPEs-mediated early polyadenylation might delay the onset of total-RNA degradation. Similar late effects are also associated with GC-rich signals resembling cCPEs (e.g., CCCC), including delayed onset of total-RNA degradation (p<8e-8), faster degradation (p<2e-3) and lower AT-ratios (p<2e-9 at 8hpf). Thus, GC-rich signals might delay the onset of total-RNA degradation independently of polyadenylation. Finally, A-rich signals within 3’UTRs (e.g., AAAAAA) are linked to lower AT-ratio both before (p<5e-12 at 1hpf) and after genome activation (p<5e-9 at 10hpf).

**Figure 5:**
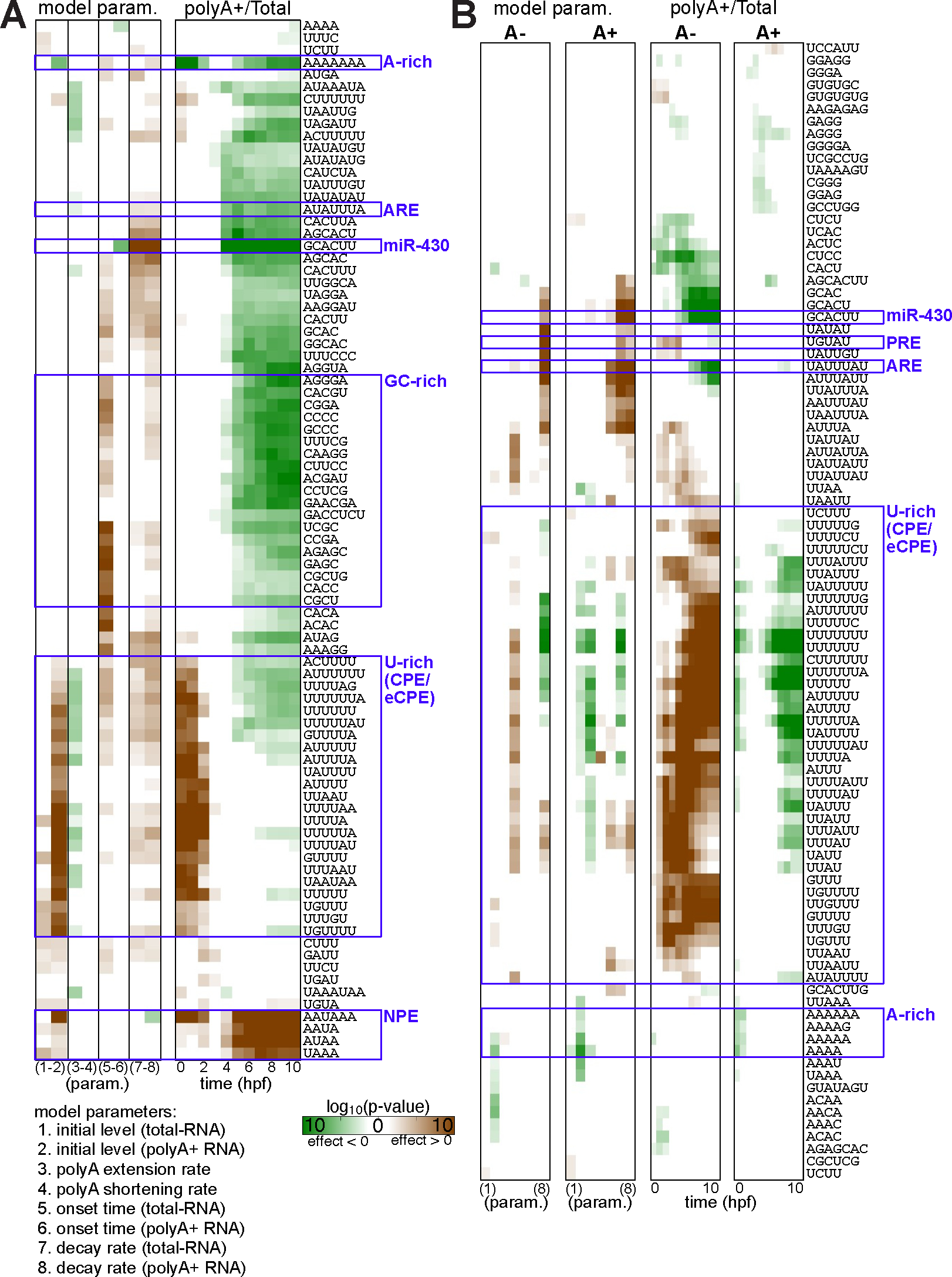
Cis-regulatory elements in 3’UTRs of maternal mRNAs **(A)** Enrichments associated with 3’UTRs of native zebrafish maternal mRNAs. Enriched k-mers (rows) in 3’UTRs relative to parameters estimated by models (columns, left) or ratio of estimated polyA+ and total RNA levels at different times (columns, right). Values represent - 1*log10(p-value) of a one-sided KS-test with 1% FDR (brown: lower mean value in instances that contain a k-mer, green: higher mean value in instances that contain a k-mer). Matrix includes the top 10 k-mers (with most significant p-values) in each column (parameter/time). List of tested parameters (by number) is indicated. De-adenylation parameter was divided into negative values (3. polyA extension rate) and positive values (4. polyA shortening rate). K-mers of specific groups are marked by blue boxes. **(B)** Enrichments associated with 3’UTRs of massively parallel mRNA reporters (A+: with an initial polyA tail of 40nt, A-: without an initial tail). Table is structured as in (A). De-adenylation parameter is only negative for A- reporters (3. polyA extension rate) and only positive for A+ reporters (4. polyA shortening rate).

Analysis of sequence elements within MPRA data (**Fig. 5B**, **Table S4**) can fine-tune the regulatory rules identified in the genomic analysis. MPRA provides enhanced statistical power and minimizes background effects by using a large set of reporters with a constant structure and a short sequence variability. However, unlike native transcripts, reporters are introduced into embryos at fixed amounts and a pre-defined initial tail length, which can affect regulation after injection. Analysis of reporters confirmed several expected degradation elements (Rabani et al., 2017). These include the destabilizing miR-430 6-mer seed (GCACUU, A+ p<1e-18), ARE (e.g., UAUUUAU, A+ p<1e-32) and Pumilio binding sites (PRE, e.g., UGUAU, A- p<6e-13); and stabilizing poly-U elements (e.g., UUUUUU, A- p<2e-16). In A- reporters, degradation effects are stronger on polyA+ RNA, while in A+ reporters the effects are stronger on total-RNA. In A- reporters, CPEs (e.g., UUUUUU) are associated with delayed onset of total-RNA degradation (p<4e-8) and higher AT-ratios (p<8e-32 at 8hpf), suggesting longer poly(A) tails, as expected. However, within A+ reporters, such CPEs are associated with slower poly(A) tail extension rates (p<8e-9) and lower AT-ratios (p<2e-18 at 8hpf), suggesting shorter poly(A) tails. In A+ reporters, A-rich signals (e.g., AAAAAA) were also linked to lower AT-ratios (p<5e-5 at 1hpf). Thus, the initial poly(A) tail of reporters indeed affects the activity of signals within their 3’UTRs.

Taken together, these results associate 3’UTR regulatory signals with the in-vivo regulation of mRNA stability and its poly(A) tail length. The effect of those signals changes between early and late phases of maternal mRNA regulation, and is also affected by the initial poly(A) tail length of injected reporters.

### Classification of embryonic genes is conserved within egg-laying vertebrates and mammalians

To generalize our zebrafish results, we applied QUANTA to analyze maternal expression patterns in embryos of additional vertebrates. We analyzed total-RNA-Seq and polyA+ RNA- Seq temporal datasets of frog embryos (Owens et al., 2016; Tan et al., 2013) during the first 14 hours of their development (end of gastrulation, as in zebrafish), and of mouse (Liu et al., 2018; Qiao et al., 2020; Wang et al., 2018; Wu et al., 2022; Xue et al., 2013) and human (Hendrickson et al., 2017; Sha et al., 2020; Wu et al., 2018; Xue et al., 2013; Yan et al., 2013) pre-implantation embryos through the morula stage (**Methods**). The western clawed frog (*Xenopus tropicalis*) embryo serves as one of the best-studied model systems of early vertebrate development. Both zebrafish and frog are egg-laying models with known similarities and differences (Briggs et al., 2018; Zhang and Sheets, 2009). On the other hand, mammalian development is more divergent, and particularly genome activation in mouse and human embryos is significantly slower (Vastenhouw et al., 2019).

Classification of embryonic genes into maternal, zygotic or combined (maternal and zygotic) expression groups changes between organisms (**Table S1**). Globally, 66%-70% of orthologs are similarly classified between organisms, while between human and mouse similarity increases to 79%. Fraction of zygotic genes remains similar in all organisms (∼10%, **Fig. 6A**). However, fraction of exclusively maternal genes is reduced from 30-35% in zebrafish and frog to 18% in human and mouse (**Fig. 6A**). Permutation tests show that classification of orthologs is more significantly conserved within egg-laying organisms (frog and zebrafish) and within mammalians (human and mouse), while conservation between these two groups is lower (**Fig. 6B**). Functional classes are also different between organisms (**Table S5**). While maternal genes in all organisms are enriched for sexual reproduction, mammalian maternal genes are also enriched for developmental processes. These differences agree with an enhanced specificity to developmental genes within maternal transcripts in mammalians, as previously observed (Shen-Orr et al., 2010). Zygotic genes in all organisms are enriched for transcription regulation and developmental processes. However, zygotic genes in egg-laying organisms are more specifically enriched for pattern specification and regionalization. In mammalians on the other hand, zygotic genes are enriched for cell proliferation. These differences reflect different timing of zygotic genome activation relative to other developmental processes, which is much earlier in mammalians. Finally, combined (maternal and zygotic) genes are enriched for housekeeping processes in all organisms, including cell cycle, translation and chromatin organization. Interestingly, DNA damage response is enriched in combined (maternal and zygotic) genes in mammalians but in exclusively maternal genes of egg-laying organisms (frog and zebrafish).

**Figure 6:**
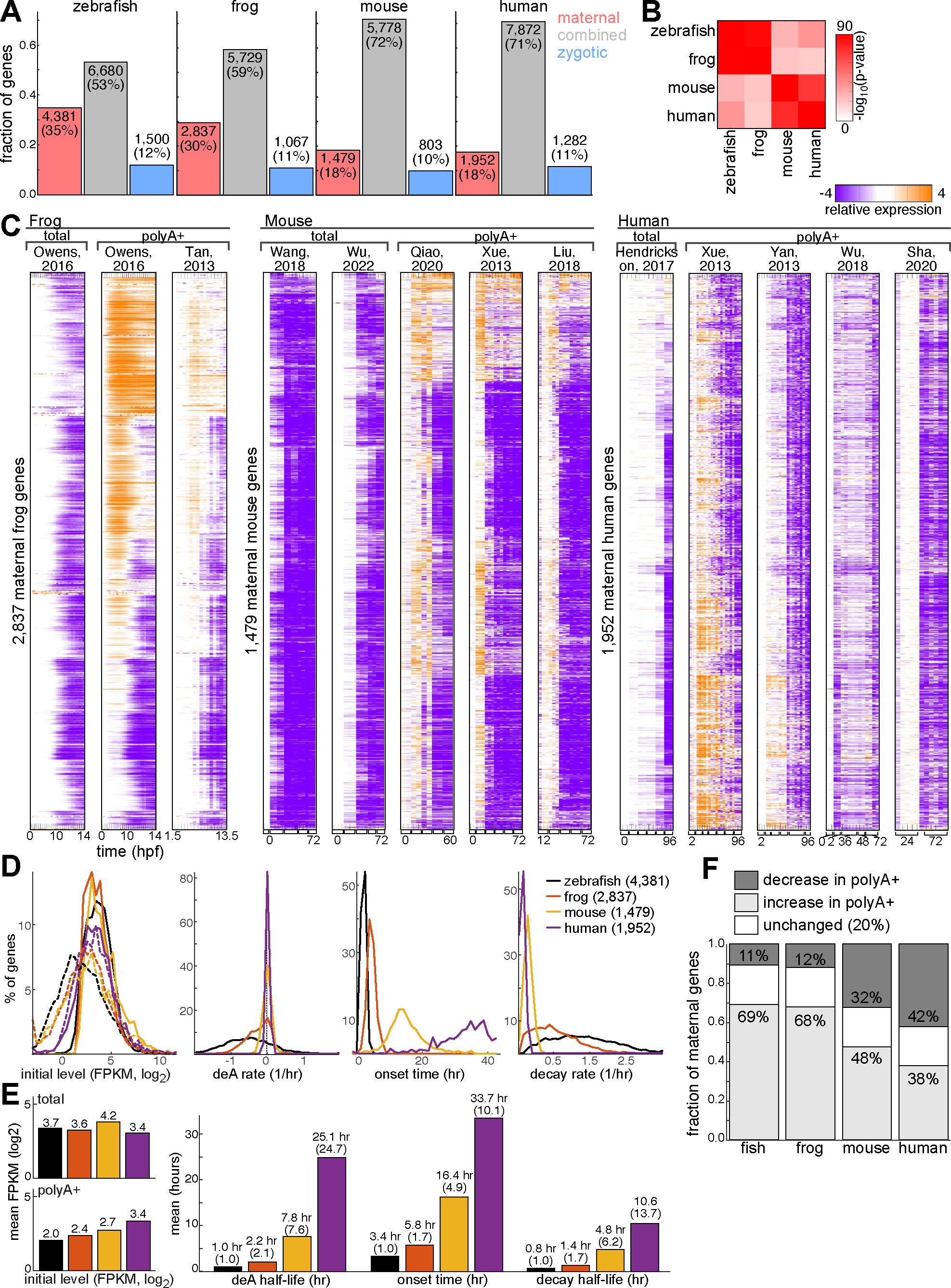
Maternal expression patterns in embryos of different organisms **(A)** Classification (x-axis: class) of embryonic genes (y-axis, fraction of genes) in four different organisms (left to right: frog, mouse and human). Number of genes and their factions are indicated on bars. **(B)** P-values for permutation tests comparing classification of embryonic genes into maternal, zygotic or combined (maternal and zygotic) expression in orthologs of 4 organisms. Color represents -log10(p-value) scale (higher = more significant). **(C)** Temporal (columns) mRNA expression patterns relative to initial measurement (orange: increase; purple: decrease; white: unchanged) of maternal genes (rows) in total-RNA-Seq datasets (left), and polyA+ RNA-Seq datasets (right) of early embryogenesis in four different organisms (left to right: frog, mouse and human). **(D)** Distribution (y-axis, % of maternal genes) of per-gene average of parameter values (x-axis) estimated by models in embryonic RNA-Seq datasets of four different organisms (red: frog, yellow: mouse, purple: human, black: zebrafish; solid: degradation model, dashed: polyA+ model). Left to right: initial RNA level (normalized FPKM, log2), deadnylation rate (1/hr), onset/switch time (hr) and decay rate (1/hr). **(E)** Fraction of maternal genes (y-axis) with either a negative deadenylation rate (increase in polyA+ fraction, light gray) or a positive deadenylation rate (decrease in polyA+ fraction, dark gray) for each organism (x-axis). A subset of 20% of genes with rates closest to zero (in absolute value) was defined as non-changing (white). **(F)** Mean values (y-axis) of regulatory parameters predicted by QUANTA in maternal genes of 4 organisms (red: frog, yellow: mouse, purple: human, black: zebrafish). Mean value (hours) and scale relative to zebrafish are indicated on bars.

### Conserved principles of maternal mRNA degradation scale by developmental pace

Next, we focused our analysis on maternal genes, and compare their expression kinetics between organisms. This comparison reveals conserved principles of maternal mRNA kinetics across organisms. In all organisms, expression levels in polyA+ fraction initially increase for many genes (**Fig. 6C**), allowing an improved fit to the ‘polyA+’ model, while total-RNA levels remain constant and retain the ‘degradation’ model (**Fig. S7**). Initial expression levels have a similar range in all organisms (**Fig. 6D-E**). PolyA+ RNA levels are 3-fold lower on average than total RNA levels, suggesting many maternal mRNAs are deposited into the embryo with short poly(A) tails. On average, 68-69% of maternal genes are polyadenylated early in zebrafish and frog (negative deadenylation rate), while this fraction is reduced to 38-48% in mouse and human (**Fig. 6F**). As in zebrafish, degradation initiates at genome activation (**Fig. 6D**), and degradation rates are correlated between polyA+ to total RNA.

Interestingly, maternal mRNA degradation kinetics is proportional to developmental pace of each organism. Onset of degradation (**Fig. 6D-E**) matches expected time of zygotic genome activation in each organism (3.5 hpf zebrafish, 4.5 hpf frog, 22 hpf mouse, 32 hpf human). In addition, both deadenylation rates and degradation rates are faster in zebrafish and frog than human and mouse, and scaled with developmental pace. For example, maternal degradation rates in frog are on average 1.7-fold slower than zebrafish (frog mean half-life 1.4 hr. compared to 0.8 hr in zebrafish, **Fig. 6E**), matching the longer time until completion of frog gastrulation (14 hpf compared to 10 hpf in zebrafish). A parallel analysis of other tissues in these organisms (**Fig. S8A**) quantified similar distributions of degradation rates in all organisms, suggesting that such marked differences in degradation rates are unique to early embryos.

Comparison of regulatory parameters between orthologs identifies significant correlations in degradation rates and initial expression levels, while onset times and deadenylation rates are not significantly correlated (**Fig. S8B)**. Degradation rates correlate within egg-laying organisms (frog and zebrafish) and within mammalians (human and mouse), but not between the two groups (**Fig. S8B-C**). Several key maternal regulators show either similar (e.g., *btg4*, *cpeb1b*, **Fig. S8D**) or different (e.g., *dazl*, **Fig. S8D**) scaled degradation rates across organisms.

Taken together, these results show that across all analyzed species, maternal mRNAs undergo changes in poly(A) tail length but remain stable prior to zygotic genome activation. Both onset and rate of degradation scale by organisms’ developmental pace. Degradation rates are conserved between orthologs, and several key regulatory genes have similarly fast degradation.

### Linking 3’UTR regulatory elements to differences in mRNA stability between organisms

We further analyzed the 3’UTR sequences of maternal mRNAs within each specie for regulatory elements that govern maternal mRNA kinetics (**Fig. S9, Table S6**).

In frog and zebrafish embryos, early signals are linked to poly(A) tail extension and delayed degradation, while late signals accelerate degradation. Zebrafish and frog share signals to accelerate degradation after genome activation, such as the miR-430 seed (GCACUU, p<5e-12 in fish, p<2e-14 in frog, **Fig. S9A**). In both species, CPEs are associated with delayed onset of total-RNA degradation and higher AT-ratios before genome activation (**Fig. S9B**), suggesting poly(A) tail extension. After genome activation, CPEs accelerate degradation and associate with lower AT-ratios. Poly-GC signals (e.g., CCGC, CCCC) are also associated with delayed onset of total-RNA degradation in both species, but their effect on AT-ratios is different. In zebrafish, these signals associate with lower AT-ratios after genome activation, while in frog they associate with higher AT-ratios in both early and later times. Indeed, in frog embryos, C-rich signals were shown to enhance early polyadenylation (Paillard et al., 2000) and translation (Xiang et al., 2024).

In mammalians, signals linked to lower AT-ratios are active both before and after genome activation, but there are no signals to accelerate degradation. Mouse 3’UTRs contain stabilizing elements, including several CPEs (e.g., UUAUUU, p<5e-6) and other uncharacterized signals (e.g., UGCAGU, p<3e-8). These signals specifically stabilize total-RNA, and are thus also linked to lower AT-ratios after genome activation. In addition, poly-GC signals in mammalian associate with delayed onset of total-RNA degradation and lower AT-ratios both before and after genome activation. Finally, A-rich elements are also associated with lower AT-ratios (e.g., AAAAAAA, frog p<3e-4, mouse p<3e-11 and human 6e-4), similar to their activity in zebrafish.

Taken together, these results suggest a different 3’UTR regulatory logic across organism.

### Massively parallel reporters demonstrate temperature dependent coupling of mRNA degradation to developmental pace

To further investigate the effect of changes in pace of development on maternal mRNA stability and the activity of 3’UTR elements, we applied UTR-Seq to zebrafish embryos grown at either 22°C or 34°C (**Fig. 7A**). Growth temperature affects the pace of zebrafish development (Urushibata et al., 2021). While in the laboratory, zebrafish embryonic development is carried out at 28°C, a higher temperature increases developmental pace, and a lower temperature slows it down. In particular, embryos reach the end of gastrulation 2 hours faster at 34°C, and 5 hours slower at 22°C (**Fig. 7B**). Overall, pace of development is 2-fold faster at 34°C compared to 22°C.

**Figure 7:**
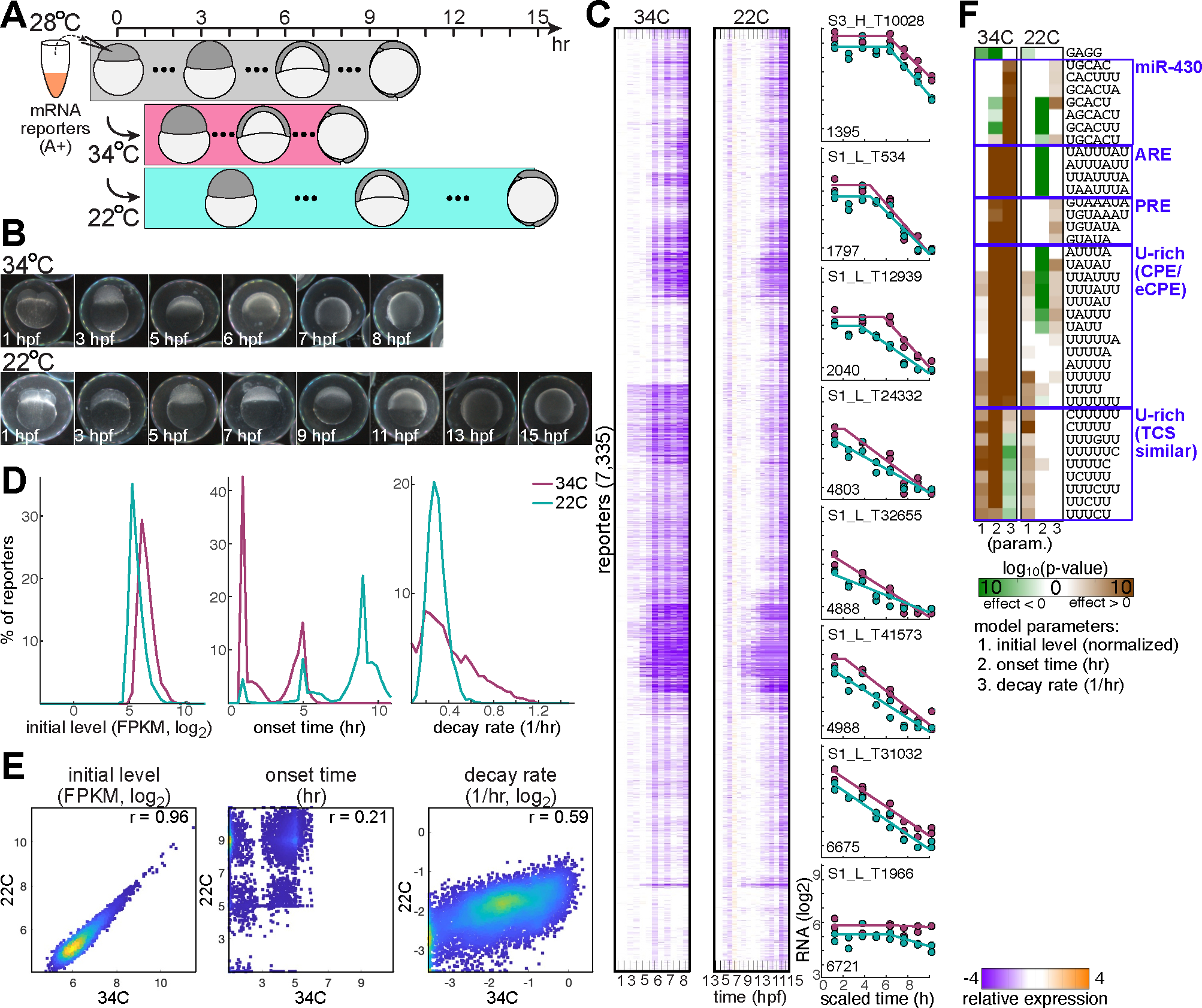
Cis-regulatory elements in 3’UTRs of maternal mRNAs across organisms **(A)** A+ UTR-Seq reporters were injected into 1-cell stage embryos collected at a standard 28C temperature. Following injection, embryos were grown at either 34C (magenta, 6C above standard temperature) or 22C (cyan, 6C below normal temperature) and followed over time. High temperature led to faster developmental progression (by 2hr). Low temperature led to slower developmental progression (by 5hr). Reporters were sequenced from temporal total RNA samples. **(B)** Embryos imaged during development at either 34C (top) or 22C (bottom). (C) Left: temporal (columns) mRNA expression patterns relative to initial measurement (orange: increase; purple: decrease; white: unchanged) of 7,335 mRNA reporters (rows) that were injected into zebrafish embryos at the 1-cell stage. Embryos were grown at a high 34C temperature (left), or at a low 22C temperature (right). Right: examples of temporal (x-axis, scaled time to match 28C growth times) expression levels (y-axis, normalized FPKM, log2) of individual reporters (magenta: 34C growth temperature, cyan: 22C growth temperature). **(D)** Distribution (y-axis, % of reporters) of parameter values (x-axis) estimated by the ‘degradation’ model in UTR-Seq datasets (magenta: 34C growth temperature, cyan: 22C growth temperature). Left to right: initial RNA level (normalized FPKM, log2), onset time (hr) and decay rate (1/hr). **(E)** Correlation of parameters (as indicated on top) estimated by the ‘degradation’ model in two UTR-Seq datasets (x-axis: 34C growth temperature, y-axis: 22C growth temperature). Pearson r value is indicated. Color represents density (blue = low density; yellow = high density). **(F)** Enriched k-mers (rows) in 3’UTRs of UTR-Seq reporters (left: 34C growth temperature, right: 22C growth temperature). Values represent -1*log10(p-value) of a one-sided KS-test with 1% FDR (brown: lower mean value in instances that contain a k-mer, green: higher mean value in instances that contain a k-mer). Matrix includes the top 10 k-mers (with most significant p-values) in each column (parameter/time). K-mers of specific groups are marked by blue boxes.

Degradation of UTR-Seq reporters scaled with developmental pace. Reporters show similar degradation profiles in 22°C or 34°C when time is scaled by developmental stages (**Fig. 7C**). Degradation kinetics is faster in 34°C, including both onset (mean onset 2.9 hpf at 34°C, 8 hpf at 22°C, **Fig. 7D**) and decay rates (median half-life 2 hpf at 34°C, 2.4 hpf at 22°C, **Fig. 7D**). Rates of degradation are correlated between the two temperatures (Pearson r=0.59, **Fig. 7E**). However, very fast degradation rates are not achieved at 22°C. Interestingly, at 34°C degradation onsets at two main times (1 hpf and 5 hpf). At 22°C there is an additional onset time at 9 hpf, but some reporters still onset in each of the earlier peaks.

Regulatory elements show distinct activities at each temperature (**Fig. 7F, Table S7**). As degradation rates at 34°C reach higher values, signals for accelerating degradation show a more significant effect at this temperature. These include the miR-430 seed (p<3*10^-61^ at 34°C, p<2*10^-8^ at 22°C), ARE (p<6*10^-117^ at 34°C) and PRE (p<1*10^-24^ at 34°C, p<6*10^-5^ at 22°C).

On the other hand, many signals that accelerate degradation at 34°C, are instead associated with earlier onset at 22°C and delayed onset at 34°C. These include AREs, PREs and U-rich CPEs. The miR-430 signal is associated with earlier onset at both 22°C and 34°C, but the effect is more significant at 22°C. These reporters are depleted from the third onset peak and enriched in the first two onset peaks at 22°C. The third peak could therefore represent very stable reporters with lower degradation, so elements that accelerate degradation are depleted from this peak. Finally, U-rich signals with a single G or C base, resembling TCS elements (Charlesworth et al., 2013), are associated with delayed onset of degradation and lower degradation rates at 34°C but not 22°C. Their stabilizing effect could be evident only in the background of enhanced degradation at 34°C.

Taken together, these results confirm the scaling of maternal mRNA degradation in the embryo with developmental pace. The activity of some regulatory signals is enhanced by developmental pace, supporting scaling of degradation kinetics and providing enhanced sensitivity for their detection.

## Discussion

In this study we report the quantification of mRNA degradation kinetic rates and the analysis of their sequence-encoded determinants. This quantification is based on QUANTA, our kinetic modeling approach. QUANTA defines the subset of pre-existing transcripts that are not transcribed during a dynamic response, by analyzing intron and exon expression of genes. It models temporal changes in total and polyA+ mRNA levels of those genes, and dissects their kinetic parameters. By applying QUANTA to temporal embryonic RNA-Seq datasets, we demonstrate key principles of maternal mRNA regulation. We further introduce a massively parallel reporter assay to measure both polyA+ and total-RNA in zebrafish embryos, and combine it with QUANTA to decode interconnected cis-regulatory logic within 3’UTRs. Finally, we use QUANTA for a comparative analysis of maternal mRNA regulation in zebrafish, frog, mouse and human embryos. We find that across all analyzed species, maternal mRNAs undergo changes in poly(A) tail length but remain stable prior to zygotic genome activation. Onset and rate of degradation scale between species by their developmental pace, and is supported by differences in the 3’UTR regulatory code. Our portal (https://rabanilab.shinyapps.io/MZT_data) offers ready access to our analysis results, as a valuable resource to investigate maternal mRNA regulation across species.

QUANTA is easily applied to any temporal total and polyA+ RNA-Seq dataset, and provides several important advantages over other existing approaches. First, it does not require any manipulations that might interfere with normal development and growth, such as transcriptional arrest (Lai et al., 2019; Viegas et al., 2022) or metabolic labeling (Lugowski et al., 2018; Rabani et al., 2011; Russo et al., 2017; Tani and Akimitsu, 2012). Second, by focusing on the subset of transcriptionally silent mRNAs, QUANTA directly estimates degradation rates during a dynamic response, rather than relying on inference to distinguish the degradation of pre-existing mRNAs from the accumulation of new transcripts. Such inference from intron-containing pre-mRNAs (Furlan et al., 2020; Lee et al., 2013) or SNPs (Harvey et al., 2013) is highly sensitive to changes in splicing rates (Rabani et al., 2014) and alternative isoforms, sequencing errors or inaccurate gene annotation. Third, it expands approaches for comparing gene expression levels between polyA+ and total-RNA fractions (Slobodin et al., 2020; Winata et al., 2018), and provides quantification of polyadenylation rates from such signals to indirectly estimate poly(A) tail lengths. Finally, QUANTA’s ability to distinguish and quantify regulatory rates greatly enhances the sensitivity to predict mRNA cis-regulatory elements from genome-wide measurements, and associate them with multiple regulatory effects. By introducing a massively parallel reporter assay to measure both polyA+ and total-RNA in zebrafish embryos, we leverage QUANTA analysis to dissect the cis-regulatory code that controls mRNA kinetics. This uncovers similarities and differences in cis-regulation between organisms, and between native mRNAs and reporters. While we identify several previously studied elements (Giraldez et al., 2006; Paillard et al., 2000; Piqué et al., 2008; Voeltz and Steitz, 1998), other signals have not been recognized before. Future work will provide more detailed investigation of their function, binding factors and developmental roles in controlling embryonic mRNA kinetics.

The similarities and differences we observed across organisms provide an opportunity to understand the evolutionary processes shaping maternal mRNA regulation. Our results associate differences in maternal mRNA stability with pace of development, in agreement with studies that also associate differences in protein stability with species-specific pace of development (Rayon et al., 2020). Temporal scaling in protein stability likely resulted from global differences affecting the tempo of cellular processes rather than variation in genetic regulatory sequences (Rayon et al., 2020). We find global degradation scaling also in zebrafish by a temperature shift, which support changes in global tempo of cellular processes as the main mechanism. However, our analysis of regulatory 3’UTR sequences suggest differences in 3’UTR signals could support differences in stability between species. These are possibly also further enhanced by global cellular processes, such as inactivation of miRNA pathways in mammalian embryos (Suh et al., 2010). Species-specific differences in the composition of maternal and zygotic transcriptomes were previously linked to differences in architecture of transcriptional regulatory regions (Shen-Orr et al., 2010). These suggest that a large fraction of maternal transcripts in egg-laying animals are expressed non-functionally. A faster degradation of maternal genes in those species would support the removal of such non-functional transcripts, to minimize their effect in the embryo, while the more specific mammalian maternal transcriptome might not require very fast degradation. Finally, we observe that the effect of some signals, such as miR-430, is limited to degradation only, while others show a combined effect on both early polyadenylation and subsequent decay. These could provide a mechanism to coordinate the need to maintain maternal mRNAs stable for extended time periods in oocytes, with their subsequent destabilization in embryos upon fertilization. Applying QUANTA also during oocyte maturation could allow to elucidate and compare the mechanisms underlying both long-term transcript stabilization in oocytes and rapid transcript decay during embryogenesis. While regulatory 3’UTR sequences suggest some mechanistic insights, further work will define mechanistic details. For example, these effects could be mediated by CPEBs which change their binding partners to promote either poly(A) tail extension or shortening (Ivshina et al., 2014).

QUANTA is broadly applicable to investigate the transcriptional shut-off component of gene expression across systems and responses. Understanding of mRNA decay has been overwhelmingly neglected compared to transcriptional control, although changes to cellular transcriptomes combine the turn-off of a subset of genes, while others are turned on (Westbrook et al., 2023). Indeed, shut-off and removal of mRNAs was also often shown critical for functionality (Choi et al., 2007; Giraldez et al., 2005; Viegas et al., 2022). QUANTA provides tools to bridge this gap, and enhance the discovery of unique roles for mRNA decay in biological responses. Expression quantitative trait loci have been shown to be 2-fold enriched in human 3ʹUTRs across tissues (Griesemer et al., 2021), but only a handful of causal 3ʹUTR variants have been described. Further combining QUANTA with genetic polymorphism analysis within populations, could link phenotypic variability to differences in mRNA degradation. These will highlight critical functions in which aberrations are not tolerated, and predict cis-regulatory mechanisms that act to establish these patterns.

QUANTA results are reproducible and reliable by several tests, but could be further improved and expanded in several directions. In particular, QUANTA analysis focuses on transcriptionally silent genes, which could be relatively few in some responses, and not provide an adequate sample for broad degradation regulation. Unlike maternal mRNAs that are not transcribed prior to genome activation, in other dynamic responses or drug-induced transcriptional shutoff, genes have been actively transcribed prior to initiation of the response. Therefore, residual transcription could persist during the early timepoints and affect estimation. Additionally, average poly(A) tail lengths of maternal mRNAs are relatively short. Differences between polyA+ and total RNA fractions might be more subtle when average poly(A) tails are longer. Finally, as single-cell Total-RNA-Seq methods are starting to emerge (Salmen et al., 2022), the principles implemented in QUANTA could be combined with pseudo-time analysis to apply to scRNA-Seq datasets.

Our work provides a general strategy to quantify mRNA degradation and polyadenylation kinetics, and investigate its sequence-based rules across species and biological systems.

## Methods

### Early embryonic RNA-Seq datasets processing and expression analysis

#### Data

Early embryonic RNA-seq datasets used in this study (**Table S8**) were taken from published works involving wild-type zebrafish, wild-type clawed frog, wild-type mouse and human samples.

#### Mapping

The following reference genomes and annotations were used for alignment: zebrafish genome GRCz11 (GenBank:GCA_000002035.4) and Ensembl (Hunt et al., 2018) annotations release 102; clawed frog genome UCB_Xtro_10.0 (xt10) and Xenbase (Fisher et al., 2023) annotations XENTR_10.0; mouse genome GRCm39 and Ensembl annotations release 108; human genome GRCh38 and Ensembl annotations release 110. RNA-Seq reads were aligned to the relevant reference genome using STAR (Dobin et al., 2013) with default parameters.

#### Expression

For all organisms except frog, gene annotations were filtered to include only genes with a ’Protein stable ID’ value or ’Gene type’ equals to ’lncRNA’. Genes in frog dataset were not filtered. Organism specific annotation files were supplemented to include a single precursor mRNA for each gene. Precursors span the entire gene locus, from the earliest annotated isoform start point to the last annotated isoform end point. Isoform specific FPKM values were calculated using Cufflinks (Trapnell et al., 2012). Mature mRNA expression was calculated by summing the FPKM values of all mature mRNA isoforms. FPKM values were log2 transformed, and values below -3 were floored to this value.

#### Temporal normalization

Temporal FPKM values (log2 transformed) in each dataset were normalized by iteratively selecting a set of control genes, and normalizing all temporal samples by their mean expression. To select control genes, we fitted a linear regression model to log2- transformed means and standard deviations of genes, and select genes with a standard deviation that is lower by 2-fold or more than expected by this model. We repeated the normalization multiple times, as long as the number of selected controls increased.

#### Selecting expressed genes

For each organism, genes were selected for further analysis by the following criteria: log2(normalized FPKM)>2 in at least one temporal sample, in more than 50% of ribosomal depleted datasets. For human analysis, all datasets (ribosomal depleted and polyA selected) were used for gene selection.

### Classification of genes

A gene is defined as maternally expressed if its maximal mature-RNA expression before genome activation is larger than a threshold value, defined by 2 standard deviations below the mean (excluding values of -3 that skewed the distribution). A gene is defined as zygotically expressed by fitting at least one of the following two criteria. (1) Ratio of maximal precursor- RNA expression after genome activation to maximal precursor-RNA expression before genome activation is larger than log2(1.25), and maximal precursor-RNA expression after genome activation is larger than -2.5. (2) Ratio of mean mature-RNA expression after genome activation to mean mature-RNA expression before genome activation is larger than log2(0.75), and maximal mature-RNA expression after genome activation is larger than -2.5. Genes that match both maternal and zygotic criteria are defined as combined maternal and zygotic genes. Genome activation times per organism were defined as follows: zebrafish 3.5 hpf, frog 4.5 hpf, mouse 22 hpf and human 32 hpf. In all organisms except human, each of the ribosomal depleted dataset was classified separately, and majority vote classification was assigned for each gene. Due to inherent noise in human measurements (with highly variable genetic background), all datasets (ribosomal depleted and polyA selected) were used for classification to increase robustness.

### Degradation models

#### Degradation model

Model is defined using the following parameters: initial expression level (*X*_0_), degradation onset time (*t*_0_), degradation rate (*β*) and degradation shut-off time (*t*_1_). Equations are:

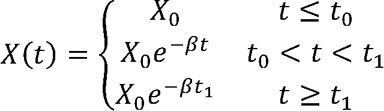

#### polyA+ model

Model is defined using the following parameters: initial expression level (*X*_0_), degradation onset time (*t*_0_), polyadenylation rate (*a*), degradation rate (*β*) and degradation shut- off time (*t*_1_). Equations are:

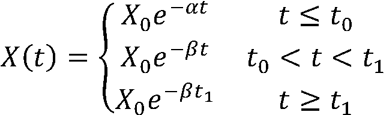

For each gene, each model was fitted to temporal RNA-Seq data using non-linear least squares regression with multiple start values (Matlab implementation). Samples with expression < 1/16*(mean of later samples) are considered dropouts and removed from the fit. The range of parameters was restricted by boundary conditions for each parameter. Initial expression level was bounded in the range (-9, 15). Onset time was bounded per organism (min: initial measurement time, max: zebrafish 6hpf, frog 9hpf, mouse 36hpf, human 48hpf). Offset time was bounded per organism (min: zebrafish 6hpf, frog 9hpf, mouse 36hpf, human 73hpf, max: latest measurement time + 1). Degradation rate was bounded to limit the range of half-life (max: zebrafish 6h, frog 9h, mouse 36h, human 73h, min: 3% of max.). Deadenylation rate was bounded to limit the range of half-life (plus/minus: zebrafish 11 min, frog 16 min, mouse 1h, human 2.2h). For prediction in UTR-Seq reporters data, we excluded the shut-off parameter (*t*_1_) to minimize over-fitting.

### Analysis of model accuracy by likelihood ratio test

We compared the fits to the two nested models (degradation model is nested in polyA+ model) by a likelihood ration test for each gene, and included an additional null model with a single parameter (constant expression level). The likelihood was calculated by assuming an additive gaussian noise model (μ*=0,* -=0.2).

### Analysis of model fitting by simulation studies

Simulated gene expression data was generated by selecting 5 values within the range of parameters fitted on each of the 7 zebrafish datasets by the two models, and applying the model to generate RNA expression data for each combination of the selected parameter values. RNA expression data was sampled according to the sampling strategy of each of the 7 zebrafish datasets. 3 total-RNA-Seq datasets were simulated using the “degradation model”, and 4 polyA+ RNA-Seq datasets were simulated using the “polyA+ model”. Random normal noise was added to simulated values (mean . = 0 and variance - = 0, 0.5, 1, 1.5). Simulated data was fitted to the degradation and polyA+ models using the same procedure as described above. We performed this fitting procedure 3 times, and took the average of 3 fitted parameter values and r-squared values between the fit and the simulated data.

### Gene set enrichment analysis

Functional enrichment analysis was performed using g:Profiler R client (version e106_eg53_p16_65fcd97) with a 1% FDR multiple testing correction method.

### Permutation tests

Classification of genes into maternal, zygotic or combined expression was compared by permutation tests. One classification was randomly permuted 1000 times relative to the other, and the number of inconsistent classifications was counted. Counts were normally distributed, and p-value was assigned using a normal distribution with mean and standard deviation as measured in permutation tests.

### Sequence k-mer enrichment analysis

For each organism, 3’UTR sequences were downloaded from the UCSC genome browser (Refseq curated and gencode for Human and Mouse) and from ensembl biomart (Yates et al., 2020). For each gene, the longest annotated 3’UTR was selected from Refseq. For genes that were not annotated in Refseq data, the longest annotated 3’UTR was selected from ensembl data. 3’UTR sequences shorter than 10bp were filtered out. Sequences were represented by a set of all short sequences between 3-7 nucleotides long (k-mers). We associated a k-mer with a regulatory effect when genes with this k-mer in their 3’UTR had a significantly different distribution (one-sided Kolmogorov-Smirnov test, 1% FDR) of a specific parameter than genes without this sequence. We assigned an effect size to each k-mer by calculating the standardized mean difference defined as 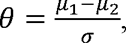, where μ_2_ is the mean of the first population, σ is the mean of the second population and - is the standard deviation (based on both populations). Within each sample, significantly enriched k-mers were filtered based on p-value and sequence similarity, using the following rule: filter out k-mers with a lower p-value that are either (1) fully contained within another k-mer or (2) fully contain another k-mer with a lower (more significant) p-value.

### Comparison between the organisms

Orthologies were calculated relative to human and relative to zebrafish. All orthology information was downloaded from ensembl biomart (Yates et al., 2020). In addition, all genes with identical gene names were considered as orthologs. Orthology information relative to zebrafish was also downloaded from the following resources. Frog orthologies were downloaded from Xenbase (Fisher et al., 2023). Mouse orthologies were downloaded from zfin (Bradford et al., 2022) and from (Giffen et al., 2019). Human orthologies were downloaded from zfin (Bradford et al., 2022).

### Web portal database for comparative maternal mRNA degradation

web portal was made using shiny65 (1.7.4 version) in R (version 4.2.2). All the 19 datasets of zebrafish, xenopus, mouse, and human were included. The models’ results for maternal genes were included.

### Zebrafish

All protocols and procedures involving zebrafish were approved by the Harvard University/Faculty of Arts and Sciences Standing Committee on the Use of Animals in Research and Teaching (IACUC; Protocol #25-08) and the Hebrew University Ethics Committee (IACUC; Protocol #NS-15859). Embryos were grown and staged according to standard procedures (Kimmel et al., 1995). Zebrafish embryos from wild-type AB/TL strains were used for all experiments.

### UTR-Seq reporter library preparation, sample collection and data processing

The UTR-Seq reporters and spike-in controls plasmids were prepared as previously described (Rabani, 2021; Rabani et al., 2017). 5 mRNAs were specifically synthesized to be used as spike-in controls, and had identical structure as A+ reporters and processed together with the sample. Controls were mixed at 2-fold decreasing amounts (highest spike-in is 16-fold higher than lowest spike-in).

### Experiment comparing total- and polyA+ reporters

UTR-Seq library preparation, injection into embryos, sequencing, data processing and normalization were all done as previously described (Rabani, 2021; Rabani et al., 2017), with the following modifications. For polyA+ sequencing of UTR-Seq reporters, total-RNA was polyA selected using the NEBNext Poly(A) mRNA Magnetic Isolation Module (NEB #E7490S). Libraries were sequenced at low-scale (approx. 2 million reads per library) on an Illumina miSeq platform with 168nt single-end reads.

### Temperature specific experiments

UTR-Seq plasmid library was linearized by enzymatic restriction at the end of its 3’UTR after (A+ library) the poly(A) sequence. HiScribe® SP6 RNA Synthesis Kit (NEB) was used to in-vitro transcribe a library of mRNA reporters from linearized plasmids. One-cell staged wild-type zebrafish embryos were collected at 28C, and injected with 100pg of mRNA reporters. Following injection, embryos were grown at the specified incubation temperature (22°C or 34°C). Two samples of 20 embryos each were collected at each time-point. In the 22°C incubation experiment, samples were collected every two hours between 1 hpf to 15 hpf. In the 34°C incubation experiment, samples were collected every two hours between 1 hpf to 5 hpf and every hour between 5 hpf and 8 hpf. Total RNA was isolated using TRI-reagent (Sigma- Aldrich), after adding 4pg of spike-in control mix during the initial TRIzol lysis step. Total RNA was reverse-transcribed with Maxima H minus RT (Thermo Fisher). Resulting cDNA was amplified by 19 PCR cycles using LongAmp Hot Start Taq 2X Master Mix (NEB). RT and PCR primers were used as previously described (Rabani, 2021; Rabani et al., 2017). PCR product was digested with EcoRV to eliminate empty vectors/reporters and cleaned with 1.0 volumes of AMPure beads (Agencourt). Libraries were quantified by Qubit fluorometric quantification (ABP Biosciences), and sequenced at low-scale (approx. 2 million reads per library) on Illumina NovaSeq platform with 122nt single-end reads. Sequencing data processing and normalization were all done as previously described (Rabani, 2021; Rabani et al., 2017).

## Author contribution statement

MR conceived and designed the project. MR wrote the QUANTA code and modeling analysis, based on initial versions for some parts of the code written by SH, DA and LF. MR and MT performed MPRA experiments. DA downloaded and analyzed zebrafish RNA-Seq datasets. PG downloaded and analyzed RNA-Seq datasets of all other organisms. PG and DA built the web portal. MR wrote the manuscript with input from the other authors. All authors read and approved the final manuscript.

## Acknowledgements

We thank Alex Schier for supporting early aspects of this project, for helpful discussions, and for providing access to sequencing facilities and zebrafish infrastructure. We thank Yotam Drier, Sagiv Shifman, Eran Meshorer and Michal Linial for critical reading of the manuscript. This research was supported by European Research Council Horizon 2020 (grant 852451 to MR) and by Israel National Science Foundation (grant 1176/21 to MR).

---

**Figure S1:**
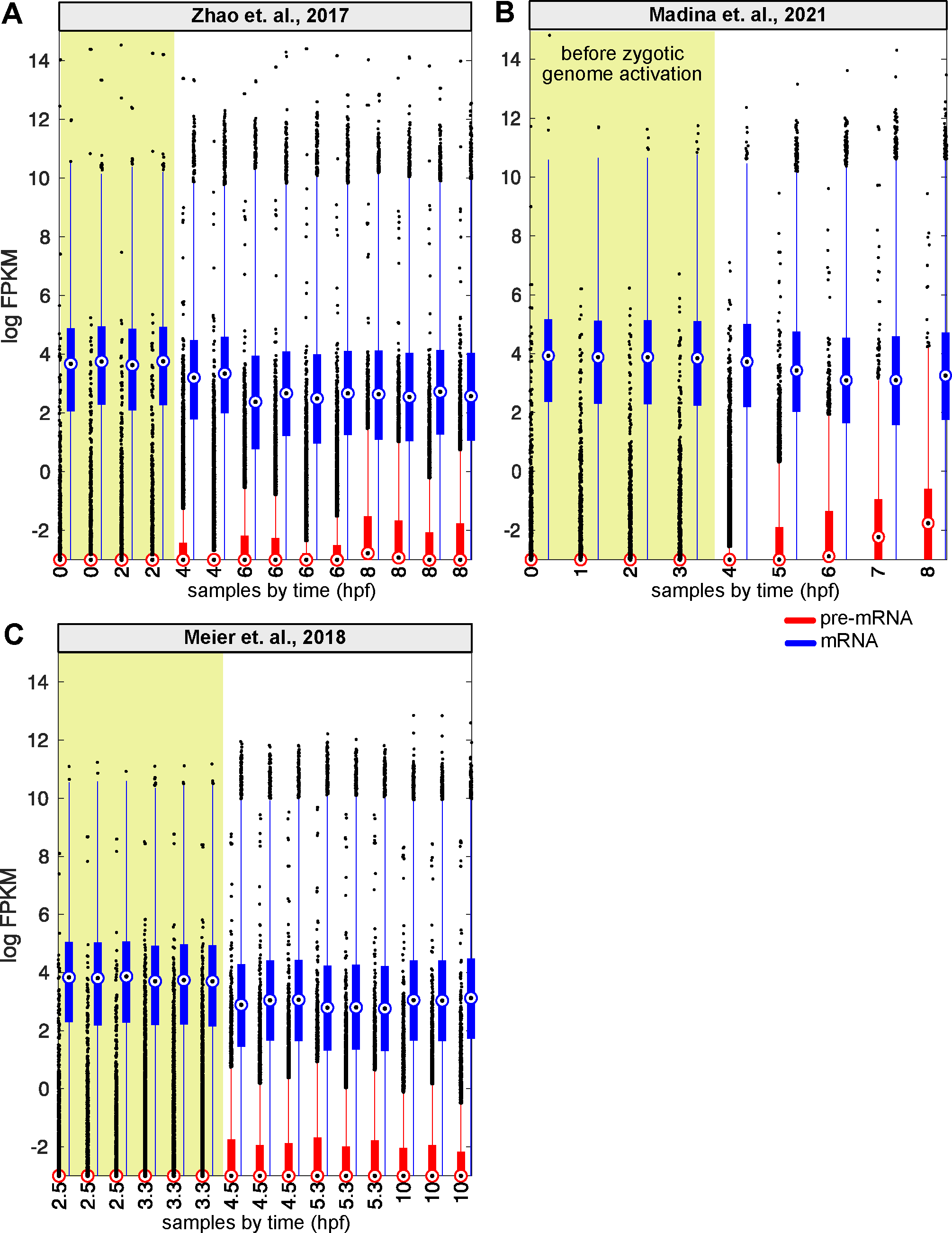
Estimated expression of pre-mRNA and mature mRNA in RNA-Seq data Distribution of normalized FPKM values (y-axis, log2) for temporal RNA-Seq samples (x- axis, hpf) in 3 zebrafish total-RNA-Seq datasets. Blue: mature mRNA. Red: pre-mRNA. Yellow background: samples corresponding to times preceding genome activation. Central dot is median; box bounds are 25th and 75th percentiles, upper and lower limits of whiskers are 1.5x interquartile ranges. Values outside of the upper and lower limits are defined as outliers **(A)** Zhao et. al., 2017. **(B)** Madina et. al., 2021. **(C)** Meier et. al., 2018.

**Figure S2:**
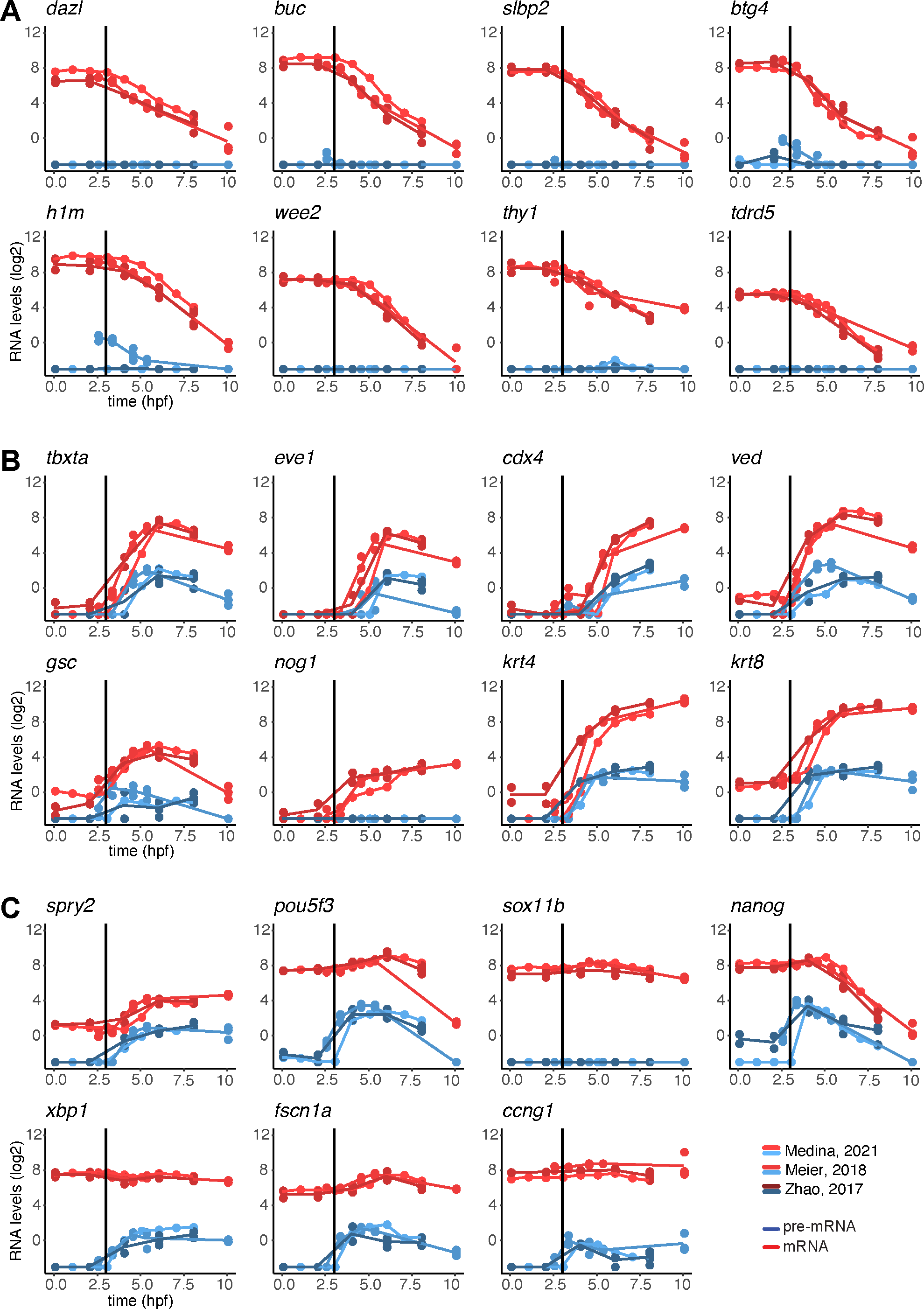
Estimated temporal expression of known embryonic genes Examples of temporal (x-axis, hpf) expression levels (y-axis, normalized FPKM, log2) measured in 3 total-RNA-Seq datasets for specific genes (names are indicated on graphs). Red: mature mRNA. Blue: pre-mRNA. **(A)** Maternal genes. **(B)** Zygotic genes. **(C)** Combined (maternal and zygotic) genes.

**Figure S3:**
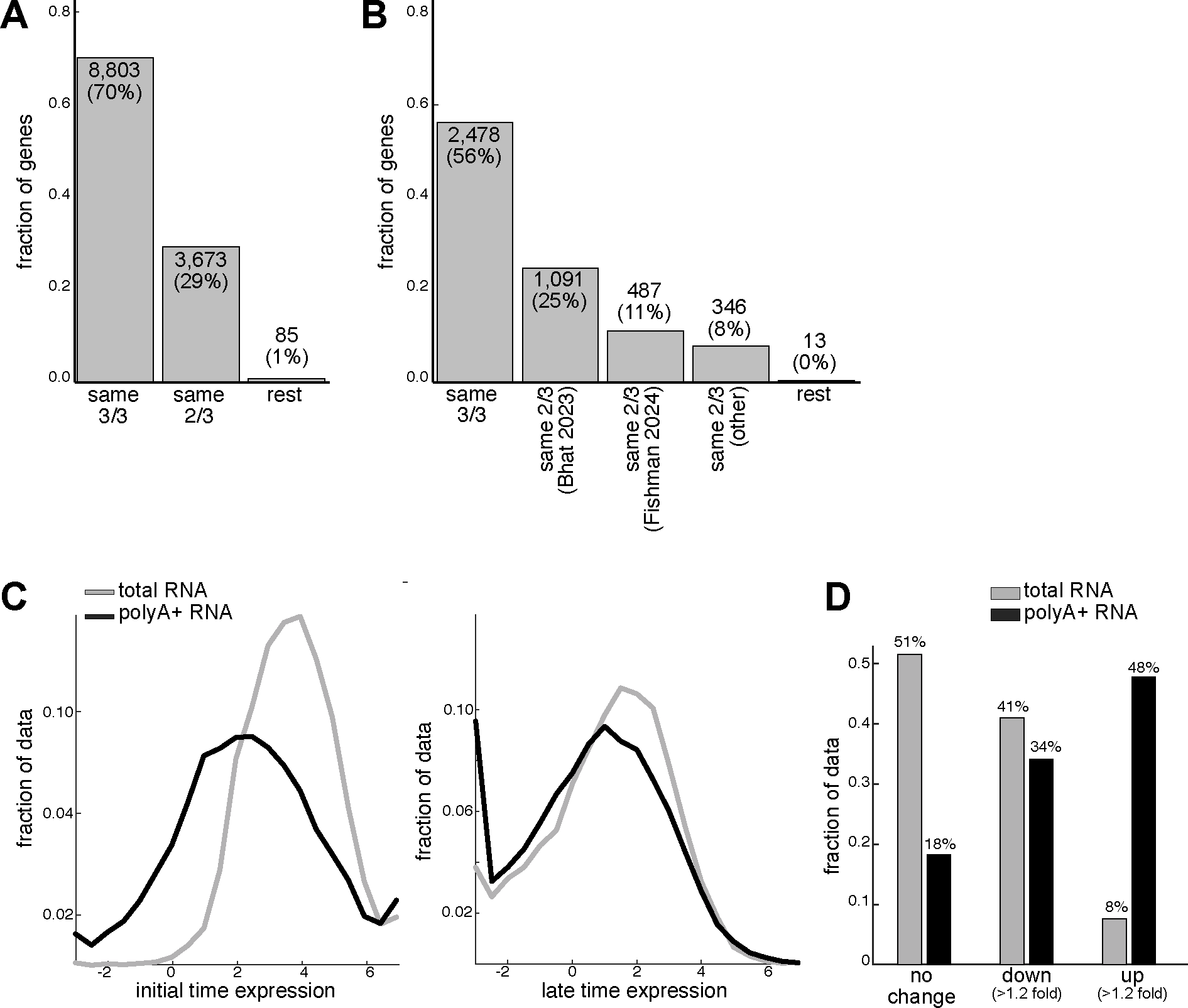
Analysis of zebrafish embryonic RNA-Seq datasets **(A)** Comparison of classification of genes into maternal, zygotic or combined genes between 3 zebrafish total-RNA-Seq embryonic genes (y-axis, fraction of genes), which were either classified similarly in all 3 datasets (3/3), in two out of 3 datasets (2/3) or differently in each datasets. Number of genes and their factions are indicated on bars. **(B)** Comparison of classification of genes into maternal, zygotic or combined genes (y-axis, fraction of genes) between classification done in this work and in two other publications based on 4sU metabolic labeling (Bhat et al., 2023; Fishman et al., 2023). Genes were either classified similarly in all 3 datasets (3/3), in two out of 3 datasets (2/3) or differently in each dataset. Number of genes and their factions are indicated on bars. **(C)** Distribution of normalized FPKM expression levels (log2, x-axis) in 3 total-RNA-Seq (gray) and 4 polyA+ RNA-Seq (black) zebrafish datasets at the initial (left) or the latest (right) measured timepoint. **(D)** Fold-change relative to the initial timepoint of normalized FPKM expression levels across 3 total-RNA-Seq (gray) and 4 polyA+ RNA-Seq (black) zebrafish datasets. Fold-change is defined as either “no change” (fold-change < 1.2), “down” (>1.2 fold reduction) or “up” (>1.2 fold increase).

**Figure S4:**
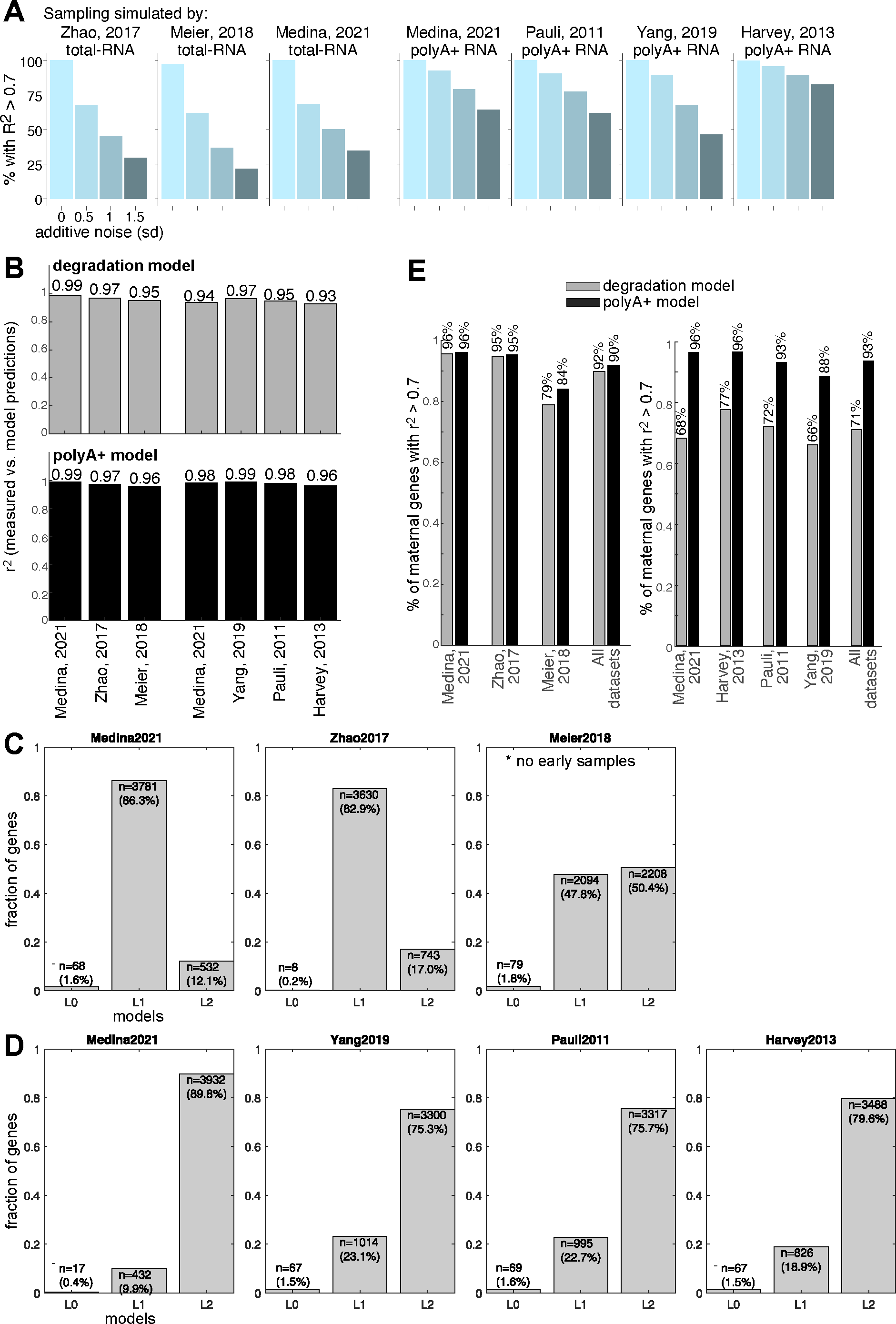
Fitting kinetic models to zebrafish embryonic RNA-Seq datasets **(A)** Results of model fitting on simulated expression datasets. Simulated gene expression data was generated by selecting 5 values within the range of parameters fitted on each of the 7 zebrafish datasets by the two models, and applying the model to generate RNA expression data for each combination of the selected parameter values. RNA expression data was sampled according to the sampling strategy of each of the 7 zebrafish datasets. 3 total-RNA-Seq datasets were simulated using the “degradation model”, and 4 polyA+ RNA-Seq datasets were simulated using the “polyA+ model”. Random normal noise was added to simulated values (mean ! = 0 and variance $ = 0, 0.5, 1, 1.5, y-axis). Simulated data was fitted to the degradation and polyA+ models 3 times, and r-squared value between the fit and the simulated data was calculated (y-axis: fraction of simulated genes with R-squared > 0.7). **(B)** R-squared values comparing degradation (top, gray) and polyA+ (bottom, black) model predictions with measurements (y-axis) in each of 3 total-RNA-Seq and 4 polyA+ RNA-Seq zebrafish datasets (x-axis) across all 4,381 maternal genes and timepoints. R-squared values are also noted on bars. **(C)** Fraction of maternal genes (y-axis) associated with each of 3 nested models (L0, constant model; L1, “degradation” model and L2 “polyA+” model, x-axis) by likelihood ratio test in 3 total-RNA-Seq zebrafish datasets (noted on top). Number of genes and their percentage are noted on bars. **(D)** Fraction of maternal genes (y-axis) associated with each of 3 nested models (L0, constant model; L1, “degradation” model and L2 “polyA+” model, x-axis) by likelihood ratio test in 4 polyA+ RNA-Seq zebrafish datasets (noted on top). Number of genes and their percentage are noted on bars. **(E)** Fraction of maternal genes with R-squared > 0.7 (y-axis) for comparison of degradation (gray) or polyA+ (black) model predictions with measurements in each of 3 total-RNA-Seq (left) and 4 polyA+ RNA-Seq (right) zebrafish datasets (x-axis), or in the combination of all datasets together.

**Figure S5:**
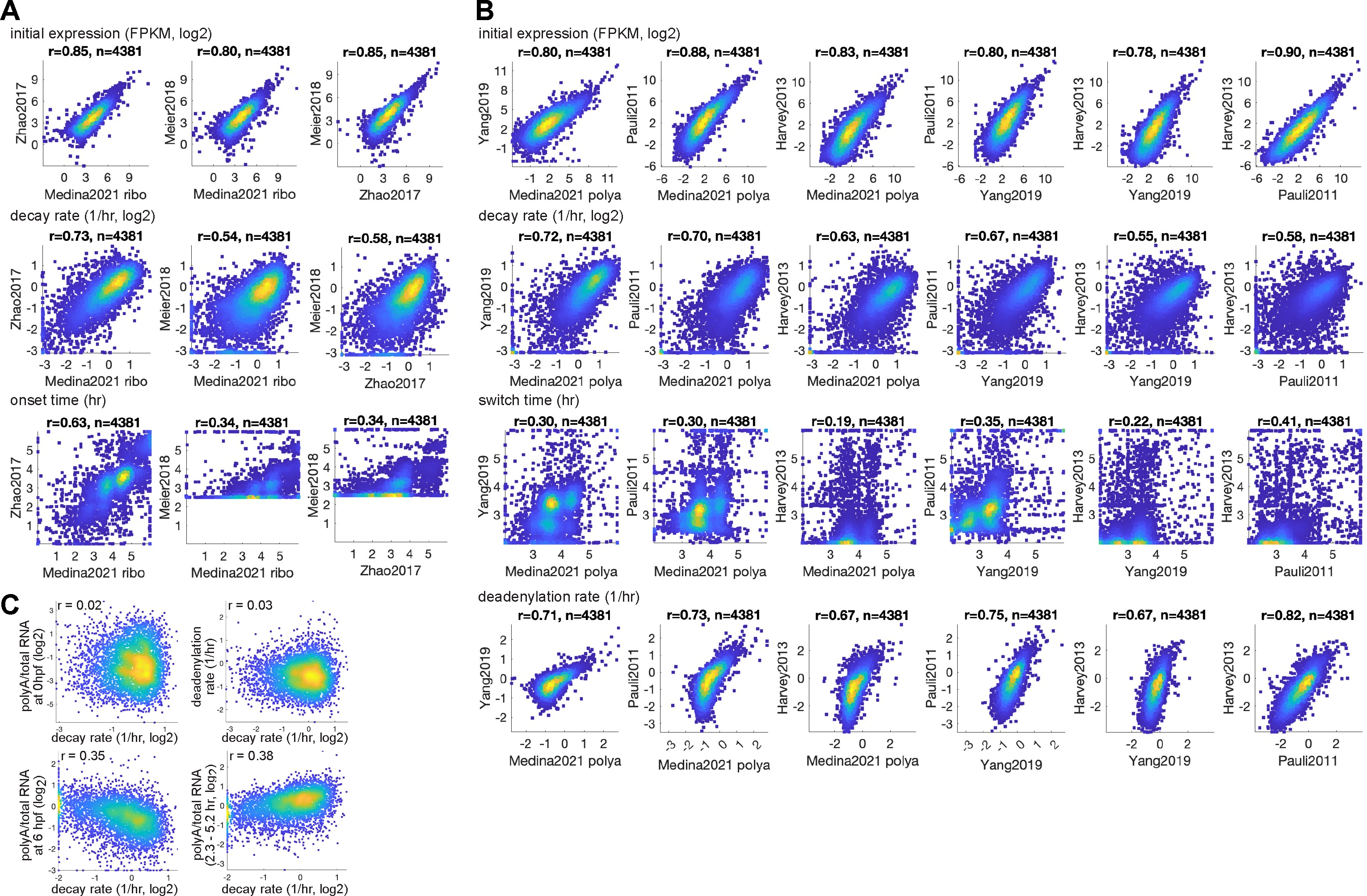
Comparison of kinetic parameters estimated in different embryonic RNA-Seq datasets **(A)** Correlation between parameter values predicted by ‘degradation model’ on 3 total-RNA- Seq zebrafish datasets. Top to bottom: initial expression level (normalized FPKM, log2), degradation rate (1/hr, log2) and onset time (hr). **(B)** Correlation between average parameter values predicted by ‘polyA+ model’ on 4 polyA+ RNA-Seq zebrafish datasets. Top to bottom: initial expression level (normalized FPKM, log2), degradation rate (1/hr), switch time (hr) and polyadenylation rate (1/hr, log2). Pearson r value and number of genes is indicated. Color represents density (blue = low density; yellow = high density). **(C)** Correlations between degradation rate (x-axis, log2) and several values estimated by our models. Top-left: the ratio of estimated polyA+ and total RNA levels at 0 hpf (y-axis). Top-right: deadenylation rates (y- axis). Bottom-left: the ratio of estimated polyA+ and total RNA levels at 6 hpf (y-axis). Bottom-right: the difference in ratio of estimated polyA+ and total RNA levels at 2.3 and 5.2 hpf (y-axis). Pearson r value is indicated (top). Color represents density (blue = low density; yellow = high density).

**Figure S6:**
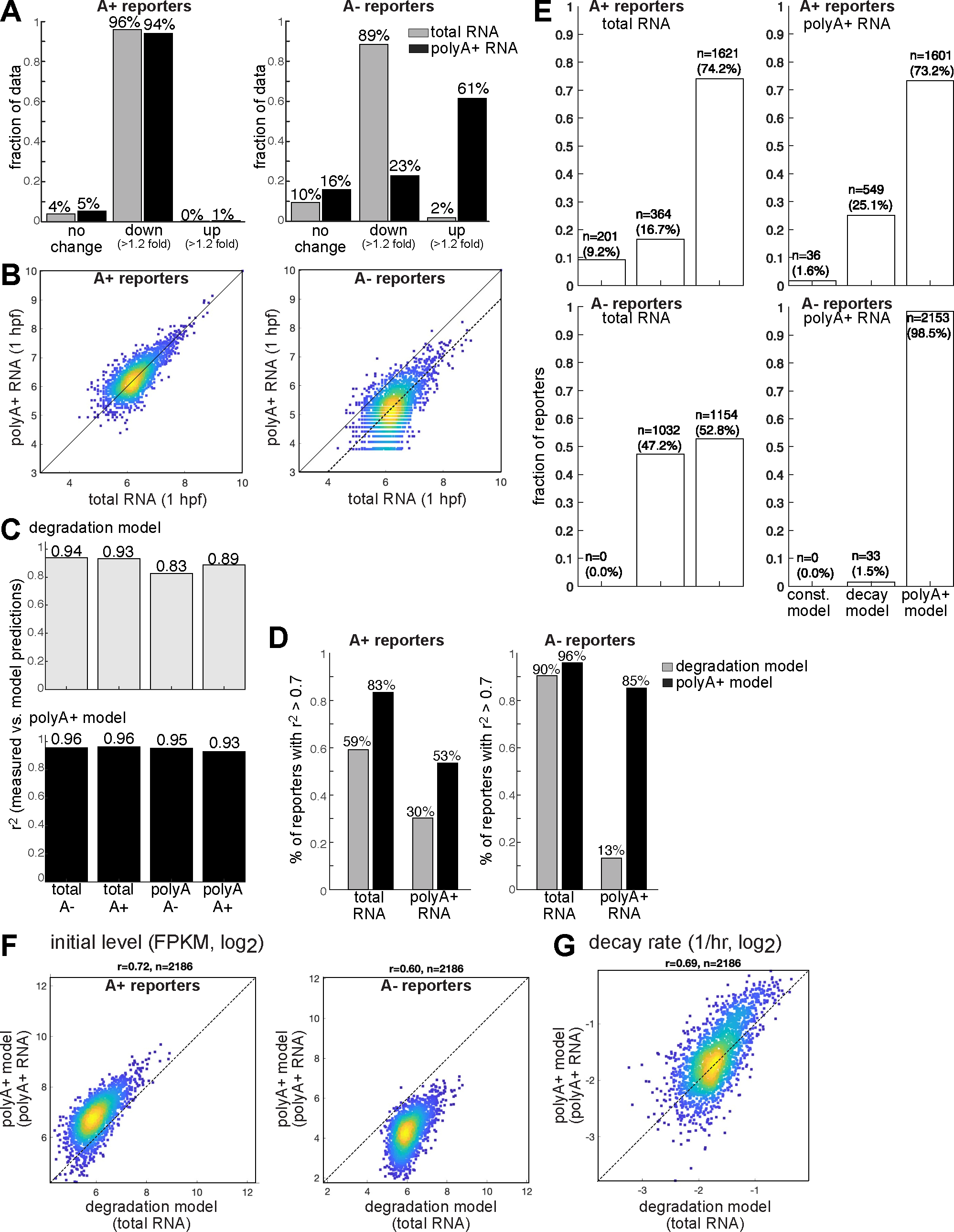
Analysis of massively parallel mRNA reporters’ expression in zebrafish embryos **(A)** Fold-change relative to the initial timepoint of normalized FPKM expression levels across total-RNA-Seq (gray) and polyA+ RNA-Seq (black) of UTR-Seq reporters’ datasets (left: A+ reporters, right: A- reporters). Fold-change is defined as either “no change” (fold-change < 1.2), “down” (>1.2 fold reduction) or “up” (>1.2 fold increase). **(B)** Correlation between normalized FPKM expression levels in total (x-axis) and polyA+ (y-axis) UTR-Seq reporters’ data (left: A+ reporters, right: A- reporters). Color represents density (blue = low density; yellow = high density). **(C)** R-squared values comparing degradation (top, gray) and polyA+ (bottom, black) model predictions with measurements (y-axis) in each of 4 UTR-Seq reporters’ datasets (x-axis). R-squared values are also noted on bars. **(D)** Fraction of maternal genes with R-squared > 0.7 (y-axis) for comparison of degradation (gray) or polyA+ (black) model predictions with measurements of UTR-Seq reporters (left: A+ reporters, right: A- reporters) by total or polyA+ RNA-Seq (x-axis). **(E)** Fraction of reporters (y-axis) associated with each of 3 nested models (constant model; “degradation” model and “polyA+” model, x-axis) by likelihood ratio test in total-RNA-Seq or polyA+ RNA-Seq datasets (noted on top). Number of genes and their percentage are noted on bars. **(F)** Correlation of initial expression levels (normalized FPKM, log2) predicted by ‘degradation model’ on total-RNA-Seq (x-axis) and ‘polyA+ model’ on polyA+ RNA-Seq (y-axis) of UTR-Seq reporters’ datasets (left: A+ reporters, right: A- reporters). Color represents density (blue = low density; yellow = high density). Number of reporters and Pearson r values are indicated. **(G)** Correlation of average decay rates (1/hr, log2) predicted by ‘degradation model’ on total-RNA-Seq (x-axis) and by ‘polyA+ model’ on polyA+ RNA-Seq (y-axis) of UTR-Seq reporters’ datasets. Rates are average of A+ and A- reporters’ predictions. Color represents density (blue = low density; yellow = high density). Number of reporters and Pearson r values are indicated.

**Figure S7:**
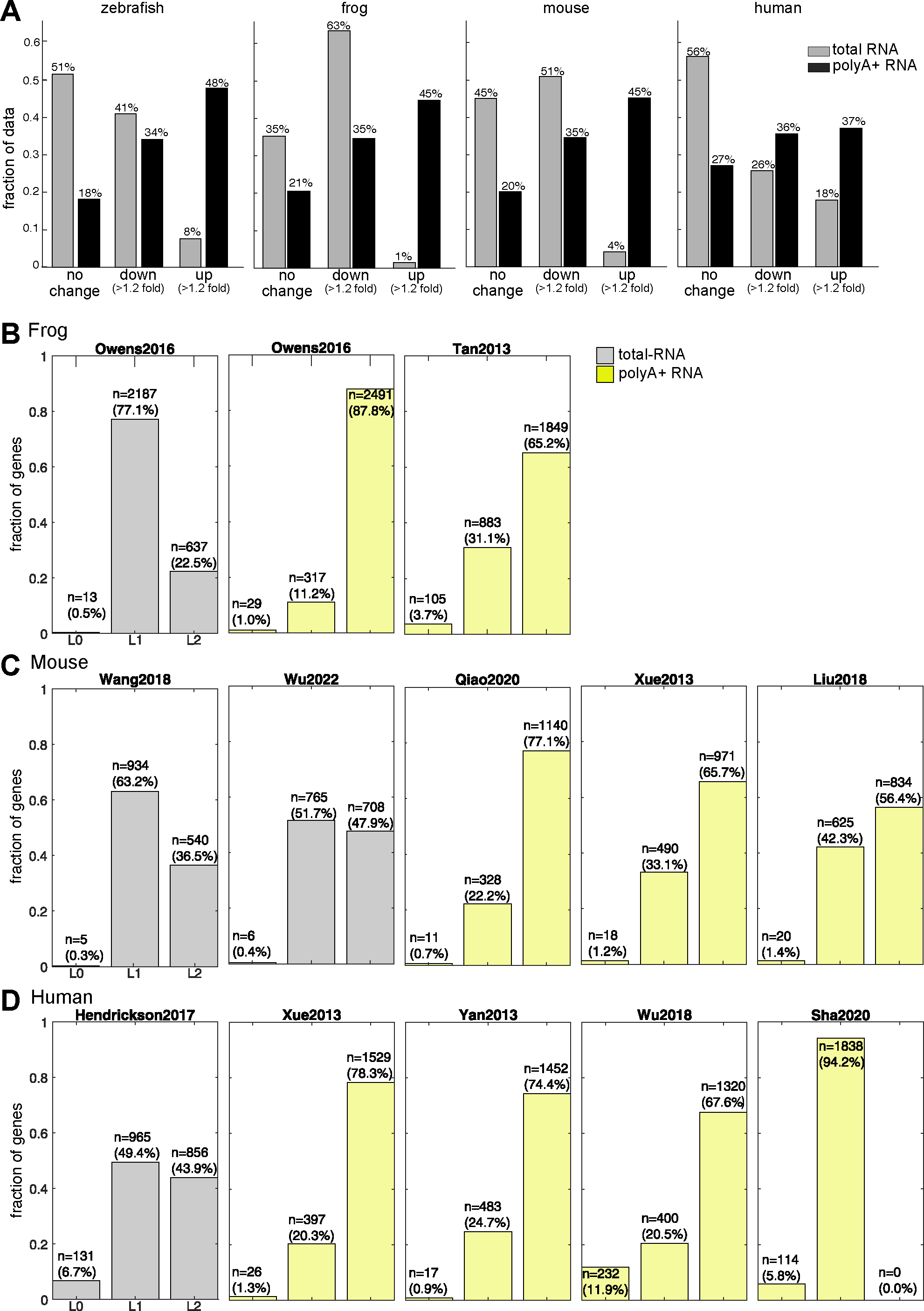
Fitting kinetic models to embryonic RNA-Seq datasets of different organisms **(A)** Fold-change relative to the initial timepoint of normalized FPKM expression levels across total-RNA-Seq (gray) and polyA+ RNA-Seq (black) datasets in four organisms (left to right: zebrafish, frog, mouse and human). Fold-change is defined as either “no change” (fold-change < 1.2), “down” (>1.2 fold reduction) or “up” (>1.2 fold increase). **(B-D)** Fraction of maternal genes (y-axis) associated with each of 3 nested models (L0, constant model; L1, “degradation” model and L2 “polyA+” model, x-axis) by likelihood ratio test in total-RNA-Seq (gray) or polyA+ RNA-Seq (yellow) datasets (noted on top). Number of genes and their percentage are noted on bars. **(B)** Frog datasets. **(C)** Mouse datasets. **(D)** Human datasets.

**Figure S8:**
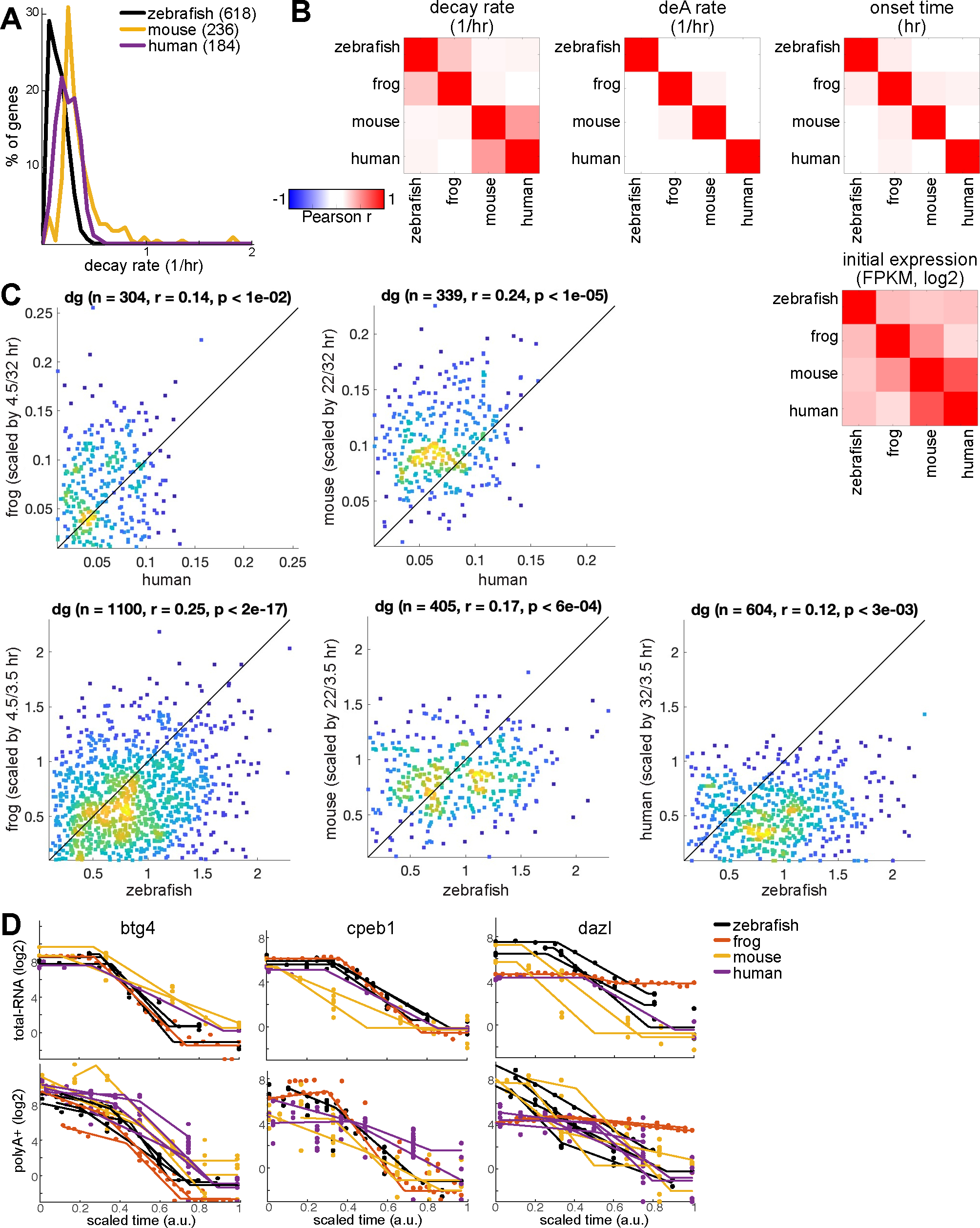
Comparative analysis of kinetic parameters across organisms **(A)** Distribution (y-axis, % of genes) of degradation rates (x-axis, 1/hr, log2) estimated for transcriptionally silent genes in total-RNA-Seq temporal data from zebrafish retina (black), mouse immune dendritic cells (yellow) and human embryonic cortex (purple). **(B)** Pearson correlation coefficient in comparison of regulatory parameters predicted by QUANTA in orthologs of 4 organisms. Color represent Pearson r values (red = positive correlation, blue = negative correlation). Correlation was calculated only between orthologs with initial expression (log2 FPKM) > 2 in both organisms. **(C)** Correlation between degradation rates (1/hr, log2) predicted in human (x-axis, top) or in zebrafish (x-axis, bottom) and those predicted on orthologs in one of 3 organisms (y-axis, left to right: frog, mouse, human). Pearson r value and number of genes is indicated. Color represents density (blue = low density; yellow = high density). **(D)** Examples of temporal (x-axis, scaled time, arbitrary units) expression levels (y- axis, normalized FPKM, log2) measured in total-RNA-Seq (top) and polyA+ RNA-Seq (bottom) datasets of four different organisms (red: frog, yellow: mouse, purple: human, black: zebrafish) for specific maternal genes (names are indicated on graphs).

**Figure S9:**
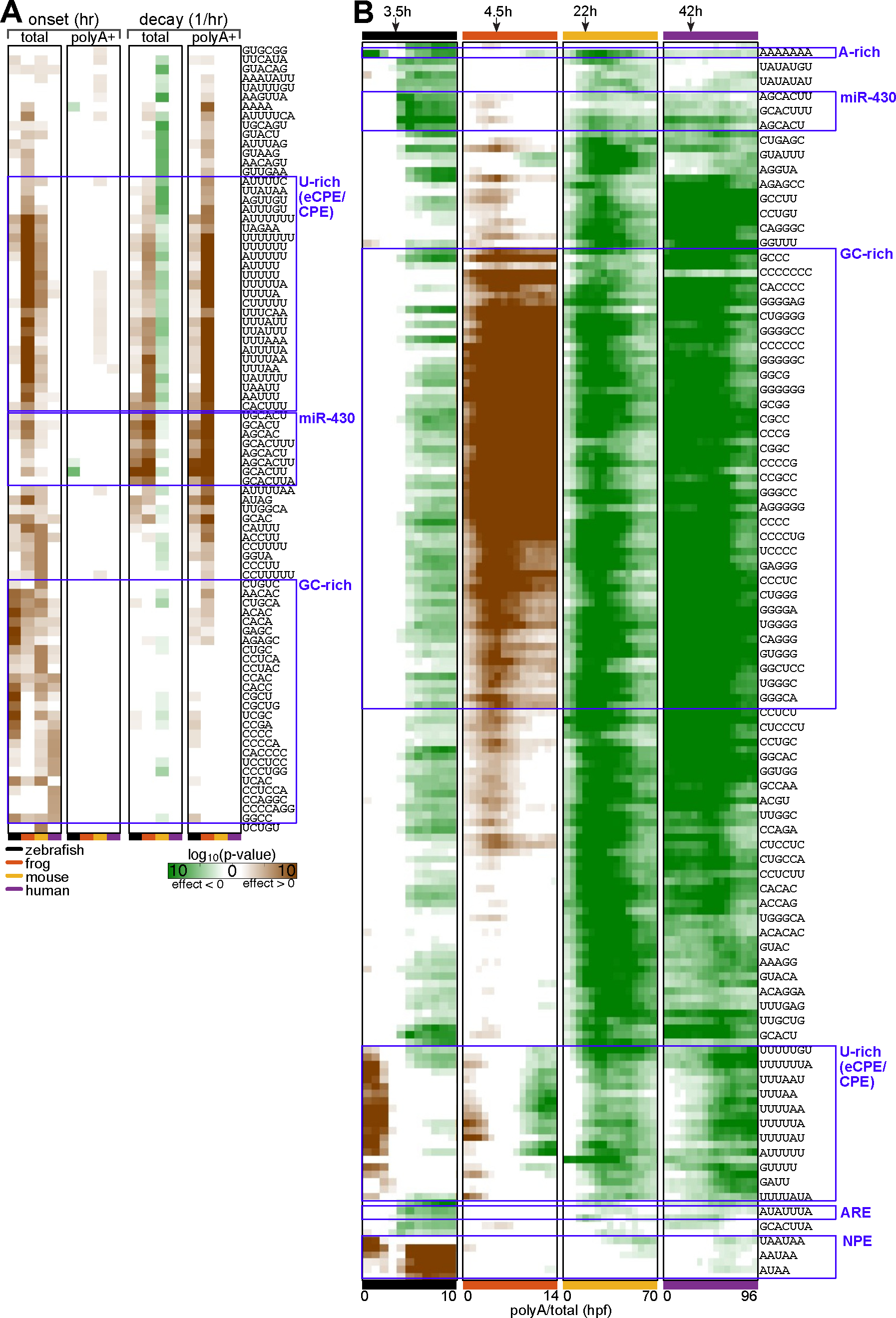
Cis-regulatory elements in 3’UTRs of maternal mRNAs across organisms Enriched k-mers (rows) in 3’UTRs of native maternal genes in four different organisms (left to right: zebrafish, frog, mouse and human). Values represent -1*log10(p-value) of a one-sided KS-test with 1% FDR (brown: lower mean value in instances that contain a k-mer, green: higher mean value in instances that contain a k-mer). Matrix includes the top 10 k-mers (with most significant p-values) in each column (parameter/time). K-mers of specific groups are marked by blue boxes. **(A)** Enrichments relative to parameters estimated by models. **(B)** Enrichments relative to estimated polyA+ and total RNA levels at different times.

## Notes

### Competing Interest Statement

The authors have declared no competing interest.

## References

1. Aanes, H., Winata, C.L., Lin, C.H., Chen, J.P., Srinivasan, K.G., Lee, S.G.P., Lim, A.Y.M., Hajan, H.S., Collas, P., Bourque, G., Gong, Z., Korzh, V., Aleström, P., Mathavan, S., 2011. Zebrafish mRNA sequencing deciphers novelties in transcriptome dynamics during maternal to zygotic transition. Genome Res 21, 1328–1338. 10.1101/gr.116012.110

2. Alonso, C.R., 2012. A complex ‘mRNA degradation code’ controls gene expression during animal development. Trends in Genetics 28, 78–88. 10.1016/j.tig.2011.10.005

3. Bailey, T.L., 2021. STREME: accurate and versatile sequence motif discovery. Bioinformatics 37, 2834–2840. 10.1093/bioinformatics/btab203

4. Bailey, T.L., Johnson, J., Grant, C.E., Noble, W.S., 2015. The MEME Suite. Nucleic Acids Res 43, W39–49. 10.1093/nar/gkv416

5. Bazzini, A.A., Del Viso, F., Moreno-Mateos, M.A., Johnstone, T.G., Vejnar, C.E., Qin, Y., Yao, J., Khokha, M.K., Giraldez, A.J., 2016. Codon identity regulates mRNA stability and translation efficiency during the maternal-to-zygotic transition. EMBO J 35, 2087–2103. 10.15252/embj.201694699

6. Bhat, P., Cabrera-Quio, L.E., Herzog, V.A., Fasching, N., Pauli, A., Ameres, S.L., 2023. SLAMseq resolves the kinetics of maternal and zygotic gene expression during early zebrafish embryogenesis. Cell Reports 42. 10.1016/j.celrep.2023.112070

7. Bradford, Y.M., Van Slyke, C.E., Ruzicka, L., Singer, A., Eagle, A., Fashena, D., Howe, D.G., Frazer, K., Martin, R., Paddock, H., Pich, C., Ramachandran, S., Westerfield, M., 2022. Zebrafish information network, the knowledgebase for Danio rerio research. Genetics 220, iyac016. 10.1093/genetics/iyac016

8. Briggs, J.A., Weinreb, C., Wagner, D.E., Megason, S., Peshkin, L., Kirschner, M.W., Klein, A.M., 2018. The dynamics of gene expression in vertebrate embryogenesis at single- cell resolution. Science 360, eaar5780. 10.1126/science.aar5780

9. Chang, H., Yeo, J., Kim, J.-G., Kim, H., Lim, J., Lee, M., Kim, H.H., Ohk, J., Jeon, H.-Y., Lee, H., Jung, H., Kim, K.-W., Kim, V.N., 2018. Terminal Uridylyltransferases Execute Programmed Clearance of Maternal Transcriptome in Vertebrate Embryos. Mol Cell 70, 72–82.e7. 10.1016/j.molcel.2018.03.004

10. Charlesworth, A., Meijer, H.A., de Moor, C.H., 2013. Specificity factors in cytoplasmic polyadenylation. Wiley Interdiscip Rev RNA 4, 437–461. 10.1002/wrna.1171

11. Choi, W.-Y., Giraldez, A.J., Schier, A.F., 2007. Target protectors reveal dampening and balancing of Nodal agonist and antagonist by miR-430. Science 318, 271–274. 10.1126/science.1147535

12. Dobin, A., Davis, C.A., Schlesinger, F., Drenkow, J., Zaleski, C., Jha, S., Batut, P., Chaisson, M., Gingeras, T.R., 2013. STAR: ultrafast universal RNA-seq aligner. Bioinformatics 29, 15–21. 10.1093/bioinformatics/bts635

13. Eichhorn, S.W., Subtelny, A.O., Kronja, I., Kwasnieski, J.C., Orr-Weaver, T.L., Bartel, D.P., 2016. mRNA poly(A)-tail changes specified by deadenylation broadly reshape translation in Drosophila oocytes and early embryos. Elife 5. 10.7554/eLife.16955

14. Fisher, M., James-Zorn, C., Ponferrada, V., Bell, A.J., Sundararaj, N., Segerdell, E., Chaturvedi, P., Bayyari, N., Chu, S., Pells, T., Lotay, V., Agalakov, S., Wang, D.Z., Arshinoff, B.I., Foley, S., Karimi, K., Vize, P.D., Zorn, A.M., 2023. Xenbase: key features and resources of the Xenopus model organism knowledgebase. Genetics 224, iyad018. 10.1093/genetics/iyad018

15. Fishman, L., Modak, A., Nechooshtan, G., Razin, T., Erhard, F., Regev, A., Farrell, J.A., Rabani, M., 2024. Cell-type-specific mRNA transcription and degradation kinetics in zebrafish embryogenesis from metabolically labeled single-cell RNA-seq. Nat Commun 15, 3104. 10.1038/s41467-024-47290-9

16. Furlan, M., Galeota, E., Gaudio, N.D., Dassi, E., Caselle, M., de Pretis, S., Pelizzola, M., 2020. Genome-wide dynamics of RNA synthesis, processing, and degradation without RNA metabolic labeling. Genome Res 30, 1492–1507. 10.1101/gr.260984.120

17. Giffen, K.P., Liu, H., Kramer, K.L., He, D.Z., 2019. Expression of Protein-Coding Gene Orthologs in Zebrafish and Mouse Inner Ear Non-sensory Supporting Cells. Front Neurosci 13, 1117. 10.3389/fnins.2019.01117

18. Giraldez, A.J., Cinalli, R.M., Glasner, M.E., Enright, A.J., Thomson, J.M., Baskerville, S., Hammond, S.M., Bartel, D.P., Schier, A.F., 2005. MicroRNAs regulate brain morphogenesis in zebrafish. Science 308, 833–838. 10.1126/science.1109020

19. Giraldez, A.J., Mishima, Y., Rihel, J., Grocock, R.J., Dongen, S.V., Inoue, K., Enright, A.J., Schier, A.F., 2006. Zebrafish MiR-430 Promotes Deadenylation and Clearance of Maternal mRNAs. Science 312, 75–79. 10.1126/science.1122689

20. Griesemer, D., Xue, J.R., Reilly, S.K., Ulirsch, J.C., Kukreja, K., Davis, J.R., Kanai, M., Yang, D.K., Butts, J.C., Guney, M.H., Luban, J., Montgomery, S.B., Finucane, H.K., Novina, C.D., Tewhey, R., Sabeti, P.C., 2021. Genome-wide functional screen of 3′UTR variants uncovers causal variants for human disease and evolution. Cell 184, 5247–5260.e19. 10.1016/j.cell.2021.08.025

21. Guo, H., Ingolia, N.T., Weissman, J.S., Bartel, D.P., 2010. Mammalian microRNAs predominantly act to decrease target mRNA levels. Nature 466, 835–840. 10.1038/nature09267

22. Guo, J., Qu, H., Chen, Y., Xia, J., 2017. The role of RNA-binding protein tristetraprolin in cancer and immunity. Med Oncol 34, 196. 10.1007/s12032-017-1055-6

23. Harvey, S.A., Sealy, I., Kettleborough, R., Fenyes, F., White, R., Stemple, D., Smith, J.C., 2013. Identification of the zebrafish maternal and paternal transcriptomes. Development 140, 2703–2710. 10.1242/dev.095091

24. Hendrickson, P.G., Doráis, J.A., Grow, E.J., Whiddon, J.L., Lim, J.-W., Wike, C.L., Weaver, B.D., Pflueger, C., Emery, B.R., Wilcox, A.L., Nix, D.A., Peterson, C.M., Tapscott, S.J., Carrell, D.T., Cairns, B.R., 2017. Conserved roles of mouse DUX and human DUX4 in activating cleavage-stage genes and MERVL/HERVL retrotransposons. Nat Genet 49, 925–934. 10.1038/ng.3844

25. Hennig, J., Sattler, M., 2015. Deciphering the protein-RNA recognition code: combining large-scale quantitative methods with structural biology. Bioessays 37, 899–908. 10.1002/bies.201500033

26. Holler, K., Neuschulz, A., Drewe-Boß, P., Mintcheva, J., Spanjaard, B., Arsiè, R., Ohler, U., Landthaler, M., Junker, J.P., 2021. Spatio-temporal mRNA tracking in the early zebrafish embryo. Nat Commun 12, 3358. 10.1038/s41467-021-23834-1

27. Hunt, S.E., McLaren, W., Gil, L., Thormann, A., Schuilenburg, H., Sheppard, D., Parton, A., Armean, I.M., Trevanion, S.J., Flicek, P., Cunningham, F., 2018. Ensembl variation resources. Database (Oxford) 2018. 10.1093/database/bay119

28. Ivshina, M., Lasko, P., Richter, J.D., 2014. Cytoplasmic polyadenylation element binding proteins in development, health, and disease. Annu Rev Cell Dev Biol 30, 393–415. 10.1146/annurev-cellbio-101011-155831

29. Jukam, D., Shariati, S.A.M., Skotheim, J.M., 2017. Zygotic Genome Activation in Vertebrates. Dev Cell 42, 316–332. 10.1016/j.devcel.2017.07.026

30. Kim, S., Wysocka, J., 2023. Deciphering the multi-scale, quantitative cis-regulatory code. Molecular Cell 83, 373–392. 10.1016/j.molcel.2022.12.032

31. Kimmel, C.B., Ballard, W.W., Kimmel, S.R., Ullmann, B., Schilling, T.F., 1995 Stages of embryonic development of the zebrafish. Developmental Dynamics 203, 253–310. 10.1002/aja.1002030302

32. Krause, M., Niazi, A.M., Labun, K., Torres Cleuren, Y.N., Müller, F.S., Valen, E., 2019. tailfindr: alignment-free poly(A) length measurement for Oxford Nanopore RNA and DNA sequencing. RNA 25,1229–1241. 10.1261/rna.071332.119

33. Lai, W.S., Arvola, R.M., Goldstrohm, A.C., Blackshear, P.J., 2019. Inhibiting transcription in cultured metazoan cells with actinomycin D to monitor mRNA turnover. Methods 155, 77–87. 10.1016/j.ymeth.2019.01.003

34. Lee, M.T., Bonneau, A.R., Takacs, C.M., Bazzini, A.A., DiVito, K.R., Fleming, E.S., Giraldez, A.J., 2013. Nanog, Pou5f1 and SoxB1 activate zygotic gene expression during the maternal-to-zygotic transition. Nature 503, 360–364. 10.1038/nature12632

35. Li, B., Dewey, C.N., 2011. RSEM: accurate transcript quantification from RNA-Seq data with or without a reference genome. BMC Bioinformatics 12, 323. 10.1186/1471-2105-12-323

36. Li, Y., Yi, Y., Lv, J., Gao, X., Yu, Y., Babu, S.S., Bruno, I., Zhao, D., Xia, B., Peng, W., Zhu, J., Chen, H., Zhang, L., Cao, Q., Chen, K., 2023. Low RNA stability signifies increased post-transcriptional regulation of cell identity genes. Nucleic Acids Research 51, 6020–6038. 10.1093/nar/gkad300

37. Lim, J., Lee, M., Son, A., Chang, H., Kim, V.N., 2016. mTAIL-seq reveals dynamic poly(A) tail regulation in oocyte-to-embryo development. Genes Dev 30, 1671–1682. 10.1101/gad.284802.116

38. Liu, J., Huang, T., Chen, W., Ding, C., Zhao, T., Zhao, X., Cai, B., Zhang, Y., Li, S., Zhang, L., Xue, M., He, X., Ge, W., Zhou, C., Xu, Y., Zhang, R., 2022. Developmental mRNA m5C landscape and regulatory innovations of massive m5C modification of maternal mRNAs in animals. Nat Commun 13, 2484. 10.1038/s41467-022-30210-0

39. Liu, Y., Wu, F., Zhang, L., Wu, X., Li, D., Xin, J., Xie, J., Kong, F., Wang, W., Wu, Q., Zhang, D., Wang, R., Gao, S., Li, W., 2018. Transcriptional defects and reprogramming barriers in somatic cell nuclear reprogramming as revealed by single-embryo RNA sequencing. BMC Genomics 19, 734. 10.1186/s12864-018-5091-1

40. Lugowski, A., Nicholson, B., Rissland, O.S., 2018. DRUID: a pipeline for transcriptome-wide measurements of mRNA stability. RNA 24, 623–632. 10.1261/rna.062877.117

41. Medina-Muñoz, S.G., Kushawah, G., Castellano, L.A., Diez, M., DeVore, M.L., Salazar, M.J.B., Bazzini, A.A., 2021. Crosstalk between codon optimality and cis-regulatory elements dictates mRNA stability. Genome Biol 22, 14. 10.1186/s13059-020-02251-5

42. Meier, M., Grant, J., Dowdle, A., Thomas, A., Gerton, J., Collas, P., O’Sullivan, J.M., Horsfield, J.A., 2018. Cohesin facilitates zygotic genome activation in zebrafish. Development 145, dev156521. 10.1242/dev.156521

43. Mishima, Y., Tomari, Y., 2016. Codon Usage and 3’ UTR Length Determine Maternal mRNA Stability in Zebrafish. Mol Cell 61, 874–885. 10.1016/j.molcel.2016.02.027

44. Nicholson, A.L., Pasquinelli, A.E., 2019. Tales of Detailed Poly(A) Tails. Trends Cell Biol 29, 191–200. 10.1016/j.tcb.2018.11.002

45. Owens, N.D.L., Blitz, I.L., Lane, M.A., Patrushev, I., Overton, J.D., Gilchrist, M.J., Cho, K.W.Y., Khokha, M.K., 2016. Measuring Absolute RNA Copy Numbers at High Temporal Resolution Reveals Transcriptome Kinetics in Development. Cell Rep 14, 632–647. 10.1016/j.celrep.2015.12.050

46. Paillard, L., Maniey, D., Lachaume, P., Legagneux, V., Osborne, H.B., 2000. Identification of a C-rich element as a novel cytoplasmic polyadenylation element in Xenopus embryos. Mechanisms of Development 93, 117–125. 10.1016/S0925-4773(00)00279-3

47. Park, J.-E., Yi, H., Kim, Y., Chang, H., Kim, V.N., 2016. Regulation of Poly(A) Tail and Translation during the Somatic Cell Cycle. Mol Cell 62, 462–471. 10.1016/j.molcel.2016.04.007

48. Passmore, L.A., Coller, J., 2022. Roles of mRNA poly(A) tails in regulation of eukaryotic gene expression. Nat Rev Mol Cell Biol 23, 93–106. 10.1038/s41580-021-00417-y

49. Pauli, A., Valen, E., Lin, M.F., Garber, M., Vastenhouw, N.L., Levin, J.Z., Fan, L., Sandelin, A., Rinn, J.L., Regev, A., Schier, A.F., 2012. Systematic identification of long noncoding RNAs expressed during zebrafish embryogenesis. Genome Res 22, 577–591. 10.1101/gr.133009.111

50. Pérez-Ortín, J.E., Alepuz, P.M., Moreno, J., 2007. Genomics and gene transcription kinetics in yeast. Trends Genet 23, 250–257. 10.1016/j.tig.2007.03.006

51. Piqué, M., López, J.M., Foissac, S., Guigó, R., Méndez, R., 2008. A combinatorial code for CPE-mediated translational control. Cell 132, 434–448. 10.1016/j.cell.2007.12.038

52. Qiao, Y., Ren, C., Huang, S., Yuan, J., Liu, X., Fan, J., Lin, J., Wu, S., Chen, Q., Bo, X., Li, X., Huang, X., Liu, Z., Shu, W., 2020. High-resolution annotation of the mouse preimplantation embryo transcriptome using long-read sequencing. Nat Commun 11, 2653. 10.1038/s41467-020-16444-w

53. Rabani, M., 2021. Massively Parallel Analysis of Regulatory RNA Sequences. Methods Mol Biol 2218, 355–365. 10.1007/978-1-0716-0970-5_28

54. Rabani, M., Levin, J.Z., Fan, L., Adiconis, X., Raychowdhury, R., Garber, M., Gnirke, A., Nusbaum, C., Hacohen, N., Friedman, N., Amit, I., Regev, A., 2011. Metabolic labeling of RNA uncovers principles of RNA production and degradation dynamics in mammalian cells. Nature Biotechnology 29, 436–442. 10.1038/nbt.1861

55. Rabani, M., Pieper, L., Chew, G.-L., Schier, A.F., 2017. A Massively Parallel Reporter Assay of 3’ UTR Sequences Identifies In Vivo Rules for mRNA Degradation. Mol Cell 68, 1083–1094.e5. 10.1016/j.molcel.2017.11.014

56. Rabani, M., Raychowdhury, R., Jovanovic, M., Rooney, M., Stumpo, D.J., Pauli, A., Hacohen, N., Schier, A.F., Blackshear, P.J., Friedman, N., Amit, I., Regev, A., 2014. High-resolution sequencing and modeling identifies distinct dynamic RNA regulatory strategies. Cell 159, 1698–1710. 10.1016/j.cell.2014.11.015

57. Ramos, S.B.V., Stumpo, D.J., Kennington, E.A., Phillips, R.S., Bock, C.B., Ribeiro-Neto, F., Blackshear, P.J., 2004. The CCCH tandem zinc-finger protein Zfp36l2 is crucial for female fertility and early embryonic development. Development 131, 4883–4893. 10.1242/dev.01336

58. Ray, D., Kazan, H., Cook, K.B., Weirauch, M.T., Najafabadi, H.S., Li, X., Gueroussov, S., Albu, M., Zheng, H., Yang, A., Na, H., Irimia, M., Matzat, L.H., Dale, R.K., Smith, S.A., Yarosh, C.A., Kelly, S.M., Nabet, B., Mecenas, D., Li, W., Laishram, R.S., Qiao, M., Lipshitz, H.D., Piano, F., Corbett, A.H., Carstens, R.P., Frey, B.J., Anderson, R.A., Lynch, K.W., Penalva, L.O.F., Lei, E.P., Fraser, A.G., Blencowe, B.J., Morris, Q.D., Hughes, T.R., 2013. A compendium of RNA-binding motifs for decoding gene regulation. Nature 499, 172–177. 10.1038/nature12311

59. Rayon, T., Stamataki, D., Perez-Carrasco, R., Garcia-Perez, L., Barrington, C., Melchionda, M., Exelby, K., Lazaro, J., Tybulewicz, V.L.J., Fisher, E.M.C., Briscoe, J., 2020. Species-specific pace of development is associated with differences in protein stability. Science 369, eaba7667. 10.1126/science.aba7667

60. Russo, J., Heck, A.M., Wilusz, J., Wilusz, C.J., 2017. Metabolic labeling and recovery of nascent RNA to accurately quantify mRNA stability. Methods 120, 39–48. 10.1016/j.ymeth.2017.02.003

61. Salmen, F., De Jonghe, J., Kaminski, T.S., Alemany, A., Parada, G.E., Verity-Legg, J., Yanagida, A., Kohler, T.N., Battich, N., van den Brekel, F., Ellermann, A.L., Arias, A.M., Nichols, J., Hemberg, M., Hollfelder, F., van Oudenaarden, A., 2022. High-throughput total RNA sequencing in single cells using VASA-seq. Nat Biotechnol 40, 1780–1793. 10.1038/s41587-022-01361-8

62. Sha, Q.-Q., Zheng, W., Wu, Y.-W., Li, S., Guo, L., Zhang, S., Lin, G., Ou, X.-H., Fan, H.-Y., 2020. Dynamics and clinical relevance of maternal mRNA clearance during the oocyte-to-embryo transition in humans. Nature Communications 11, 4917. 10.1038/s41467-020-18680-6

63. Shalem, O., Dahan, O., Levo, M., Martinez, M.R., Furman, I., Segal, E., Pilpel, Y., 2008. Transient transcriptional responses to stress are generated by opposing effects of mRNA production and degradation. Molecular Systems Biology 4, 4. 10.1038/msb.2008.59

64. Shen-Orr, S.S., Pilpel, Y., Hunter, C.P., 2010. Composition and regulation of maternal and zygotic transcriptomes reflects species-specific reproductive mode. Genome Biol 11, R58. 10.1186/gb-2010-11-6-r58

65. Siddall, N.A., McLaughlin, E.A., Marriner, N.L., Hime, G.R., 2006. The RNA-binding protein Musashi is required intrinsically to maintain stem cell identity. PNAS 103, 8402–8407. 10.1073/pnas.0600906103

66. Slobodin, B., Bahat, A., Sehrawat, U., Becker-Herman, S., Zuckerman, B., Weiss, A.N., Han, R., Elkon, R., Agami, R., Ulitsky, I., Shachar, I., Dikstein, R., 2020. Transcription Dynamics Regulate Poly(A) Tails and Expression of the RNA Degradation Machinery to Balance mRNA Levels. Mol Cell 78, 434–444.e5. 10.1016/j.molcel.2020.03.022

67. Subtelny, A.O., Eichhorn, S.W., Chen, G.R., Sive, H., Bartel, D.P., 2014. Poly(A)-tail profiling reveals an embryonic switch in translational control. Nature 508, 66–71. 10.1038/nature13007

68. Suh, N., Baehner, L., Moltzahn, F., Melton, C., Shenoy, A., Chen, J., Blelloch, R., 2010. MicroRNA Function Is Globally Suppressed in Mouse Oocytes and Early Embryos. Current Biology 20, 271–277. 10.1016/j.cub.2009.12.044

69. Tadros, W., Goldman, A.L., Babak, T., Menzies, F., Vardy, L., Orr-Weaver, T., Hughes, T.R., Westwood, J.T., Smibert, C.A., Lipshitz, H.D., 2007. SMAUG is a major regulator of maternal mRNA destabilization in Drosophila and its translation is activated by the PAN GU kinase. Dev Cell 12, 143–155. 10.1016/j.devcel.2006.10.005

70. Tan, M.H., Au, K.F., Yablonovitch, A.L., Wills, A.E., Chuang, J., Baker, J.C., Wong, W.H., Li, J.B., 2013. RNA sequencing reveals a diverse and dynamic repertoire of the Xenopus tropicalis transcriptome over development. Genome Res 23, 201–216. 10.1101/gr.141424.112

71. Tani, H., Akimitsu, N., 2012. Genome-wide technology for determining RNA stability in mammalian cells: historical perspective and recent advantages based on modified nucleotide labeling. RNA Biol 9, 1233–1238. 10.4161/rna.22036

72. Trapnell, C., Roberts, A., Goff, L., Pertea, G., Kim, D., Kelley, D.R., Pimentel, H., Salzberg, S.L., Rinn, J.L., Pachter, L., 2012. Differential gene and transcript expression analysis of RNA-seq experiments with TopHat and Cufflinks. Nat Protoc 7, 562–578. 10.1038/nprot.2012.016

73. Urushibata, H., Sasaki, K., Takahashi, E., Hanada, T., Fujimoto, T., Arai, K., Yamaha, E., 2021. Control of Developmental Speed in Zebrafish Embryos Using Different Incubation Temperatures. Zebrafish 18, 316–325. 10.1089/zeb.2021.0022

74. Vastenhouw, N.L., Cao, W.X., Lipshitz, H.D., 2019. The maternal-to-zygotic transition revisited. Development 146, dev161471. 10.1242/dev.161471

75. Vejnar, C.E., Abdel Messih, M., Takacs, C.M., Yartseva, V., Oikonomou, P., Christiano, R., Stoeckius, M., Lau, S., Lee, M.T., Beaudoin, J.-D., Musaev, D., Darwich-Codore, H., Walther, T.C., Tavazoie, S., Cifuentes, D., Giraldez, A.J., 2019. Genome wide analysis of 3’ UTR sequence elements and proteins regulating mRNA stability during maternal-to-zygotic transition in zebrafish. Genome Res 29, 1100–1114. 10.1101/gr.245159.118

76. Vesterlund, L., Jiao, H., Unneberg, P., Hovatta, O., Kere, J., 2011. The zebrafish transcriptome during early development. BMC Developmental Biology 11, 30. 10.1186/1471-213X-11-30

77. Viegas, J.O., Azad, G.K., Lv, Y., Fishman, L., Paltiel, T., Pattabiraman, S., Park, J.E., Kaganovich, D., Sze, S.K., Rabani, M., Esteban, M.A., Meshorer, E., 2022. RNA degradation eliminates developmental transcripts during murine embryonic stem cell differentiation via CAPRIN1-XRN2. Dev Cell 57, 2731–2744.e5. 10.1016/j.devcel.2022.11.014

78. Viegas, J.O., Fishman, L., Meshorer, E., Rabani, M., 2023. Calculating RNA degradation rates using large-scale normalization in mouse embryonic stem cells. STAR Protoc 4, 102534. 10.1016/j.xpro.2023.102534

79. Voeltz, G.K., Steitz, J.A., 1998. AUUUA Sequences Direct mRNA Deadenylation Uncoupled from Decay during Xenopus Early Development. Mol Cell Biol 18, 7537–7545.

80. Wang, C., Liu, X., Gao, Y., Yang, L., Li, C., Liu, W., Chen, C., Kou, X., Zhao, Y., Chen, J., Wang, Y., Le, R., Wang, H., Duan, T., Zhang, Y., Gao, S., 2018. Reprogramming of H3K9me3-dependent heterochromatin during mammalian embryo development. Nat Cell Biol 20, 620–631. 10.1038/s41556-018-0093-4

81. Westbrook, E.R., Ford, H.Z., Antolović, V., Chubb, J.R., 2023. Clearing the slate: RNA turnover to enable cell state switching? Development 150, dev202084. 10.1242/dev.202084

82. Wharton, R.P., Struhl, G., 1991. RNA regulatory elements mediate control of Drosophila body pattern by the posterior morphogen nanos. Cell 67, 955–967. 10.1016/0092-8674(91)90368-9

83. White, R.J., Collins, J.E., Sealy, I.M., Wali, N., Dooley, C.M., Digby, Z., Stemple, D.L., Murphy, D.N., Billis, K., Hourlier, T., Füllgrabe, A., Davis, M.P., Enright, A.J., Busch-Nentwich, E.M., 2017. A high-resolution mRNA expression time course of embryonic development in zebrafish. Elife 6, e30860. 10.7554/eLife.30860

84. Winata, C.L., Łapiński, M., Pryszcz, L., Vaz, C., Bin Ismail, M.H., Nama, S., Hajan, H.S., Lee, S.G.P., Korzh, V., Sampath, P., Tanavde, V., Mathavan, S., 2018. Cytoplasmic polyadenylation-mediated translational control of maternal mRNAs directs maternal-to-zygotic transition. Development 145. 10.1242/dev.159566

85. Wu, J., Xu, J., Liu, B., Yao, G., Wang, P., Lin, Z., Huang, B., Wang, X., Li, T., Shi, S., Zhang, N., Duan, F., Ming, J., Zhang, X., Niu, W., Song, W., Jin, H., Guo, Y., Dai, S., Hu, L., Fang, L., Wang, Q., Li, Y., Li, W., Na, J., Xie, W., Sun, Y., 2018. Chromatin analysis in human early development reveals epigenetic transition during ZGA. Nature 557, 256–260. 10.1038/s41586-018-0080-8

86. Wu, Y., Xu, X., Qi, M., Chen, C., Li, M., Yan, R., Kou, X., Zhao, Y., Liu, W., Li, Y., Liu, X., Zhang, M., Yi, C., Liu, H., Xiang, J., Wang, H., Shen, B., Gao, Y., Gao, S., 2022. N6- methyladenosine regulates maternal RNA maintenance in oocytes and timely RNA decay during mouse maternal-to-zygotic transition. Nat Cell Biol 24, 917–927. 10.1038/s41556-022-00915-x

87. Xiang, K., Ly, J., Bartel, D.P., 2024. Control of poly(A)-tail length and translation in vertebrate oocytes and early embryos. Developmental Cell 0. 10.1016/j.devcel.2024.02.007

88. Xue, Z., Huang, K., Cai, C., Cai, L., Jiang, C., Feng, Y., Liu, Z., Zeng, Q., Cheng, L., Sun, Y.E., Liu, J., Horvath, S., Fan, G., 2013. Genetic programs in human and mouse early embryos revealed by single-cell RNA sequencing. Nature 500, 593–597. 10.1038/nature12364

89. Yan, L., Yang, M., Guo, H., Yang, L., Wu, J., Li, Rong, Liu, P., Lian, Y., Zheng, X., Yan, J., Huang, J., Li, M., Wu, X., Wen, L., Lao, K., Li, Ruiqiang, Qiao, J., Tang, F., 2013. Single-cell RNA-Seq profiling of human preimplantation embryos and embryonic stem cells. Nat Struct Mol Biol 20, 1131–1139. 10.1038/nsmb.2660

90. Yang, E., van Nimwegen, E., Zavolan, M., Rajewsky, N., Schroeder, M., Magnasco, M., Darnell, J.E., 2003. Decay rates of human mRNAs: correlation with functional characteristics and sequence attributes. Genome Res 13, 1863–1872. 10.1101/gr.1272403

91. Yang, Y., Wang, L., Han, X., Yang, W.-L., Zhang, M., Ma, H.-L., Sun, B.-F., Li, A., Xia, J., Chen, J., Heng, J., Wu, B., Chen, Y.-S., Xu, J.-W., Yang, X., Yao, H., Sun, J., Lyu, C., Wang, H.-L., Huang, Y., Sun, Y.-P., Zhao, Y.-L., Meng, A., Ma, J., Liu, F., Yang, Y.-G., 2019. RNA 5-Methylcytosine Facilitates the Maternal-to-Zygotic Transition by Preventing Maternal mRNA Decay. Mol Cell 75, 1188–1202.e11. 10.1016/j.molcel.2019.06.033

92. Yates, A.D., Achuthan, P., Akanni, W., Allen, James, Allen, Jamie, Alvarez-Jarreta, J., Amode, M.R., Armean, I.M., Azov, A.G., Bennett, R., Bhai, J., Billis, K., Boddu, S., Marugán, J.C., Cummins, C., Davidson, C., Dodiya, K., Fatima, R., Gall, A., Giron, C.G., Gil, L., Grego, T., Haggerty, L., Haskell, E., Hourlier, T., Izuogu, O.G., Janacek, S.H., Juettemann, T., Kay, M., Lavidas, I., Le, T., Lemos, D., Martinez, J.G., Maurel, T., McDowall, M., McMahon, A., Mohanan, S., Moore, B., Nuhn, M., Oheh, D.N., Parker, A., Parton, A., Patricio, M., Sakthivel, M.P., Abdul Salam, A.I., Schmitt, B.M., Schuilenburg, H., Sheppard, D., Sycheva, M., Szuba, M., Taylor, K., Thormann, A., Threadgold, G., Vullo, A., Walts, B., Winterbottom, A., Zadissa, A., Chakiachvili, M., Flint, B., Frankish, A., Hunt, S.E., IIsley, G., Kostadima, M., Langridge, N., Loveland, J.E., Martin, F.J., Morales, J., Mudge, J.M., Muffato, M., Perry, E., Ruffier, M., Trevanion, S.J., Cunningham, F., Howe, K.L., Zerbino, D.R., Flicek, P., 2020. Ensembl 2020. Nucleic Acids Res 48, D682–D688. 10.1093/nar/gkz966

93. Yu, C., Ji, S.-Y., Sha, Q.-Q., Dang, Y., Zhou, J.-J., Zhang, Y.-L., Liu, Y., Wang, Z.-W., Hu, B., Sun, Q.-Y., Sun, S.-C., Tang, F., Fan, H.-Y., 2016. BTG4 is a meiotic cell cycle– coupled maternal-zygotic-transition licensing factor in oocytes. Nat Struct Mol Biol 23, 387–394. 10.1038/nsmb.3204

94. Zhang, Y., Sheets, M.D., 2009. Analyses of zebrafish and Xenopusoocyte maturation reveal conserved and diverged features of translational regulation of maternal cyclin B1 mRNA. BMC Developmental Biology 9, 7. 10.1186/1471-213X-9-7

95. Zhao, B.S., Wang, X., Beadell, A.V., Lu, Z., Shi, H., Kuuspalu, A., Ho, R.K., He, C., 2017. m6A-dependent maternal mRNA clearance facilitates zebrafish maternal-to-zygotic transition. Nature 542, 475–478. 10.1038/nature21355

96. Zheng, W., Zhou, Z., Sha, Q., Niu, X., Sun, X., Shi, J., Zhao, L., Zhang, S., Dai, J., Cai, S., Meng, F., Hu, L., Gong, F., Li, X., Fu, J., Shi, R., Lu, G., Chen, B., Fan, H., Wang, L., Lin, G., Sang, Q., 2020. Homozygous Mutations in BTG4 Cause Zygotic Cleavage Failure and Female Infertility. The American Journal of Human Genetics 107, 24–33. 10.1016/j.ajhg.2020.05.010

